# Can MRI measure myelin? Systematic review, qualitative assessment, and meta-analysis of studies validating microstructural imaging with myelin histology

**DOI:** 10.1101/2020.09.08.286518

**Authors:** Alberto Lazari, Ilona Lipp

**Affiliations:** Wellcome Centre for Integrative Neuroimaging, FMRIB, Nuffield Department of Clinical Neurosciences, University of Oxford, UK; Department of Neurophysics, Max Planck Institute for Human Cognitive and Brain Sciences, Leipzig, Germany

**Keywords:** Myelin, MRI, validation, microstructural imaging, relaxometry, magnetization transfer, diffusion, histology

## Abstract

Recent years have seen an increased understanding of the importance of myelination in healthy brain function and neuropsychiatric diseases. Non-invasive microstructural magnetic resonance imaging (MRI) holds the potential to expand and translate these insights to basic and clinical human research, but the sensitivity and specificity of different MR markers to myelination is a subject of debate.

To consolidate current knowledge on the topic, we perform a systematic review and meta-analysis of studies that validate microstructural imaging by combining it with myelin histology.

We find meta-analytic evidence for correlations between myelin histology and markers from different MRI modalities, including fractional anisotropy, radial diffusivity, macromolecular pool, magnetization transfer ratio, susceptibility and longitudinal relaxation rate, but not mean diffusivity. Meta-analytic correlation effect sizes range widely, between *R*^2^ = 0.26 and *R*^2^ = 0.82. However, formal comparisons between MRI-based myelin markers are limited by methodological variability, inconsistent reporting and potential for publication bias, thus preventing the establishment of a single most sensitive strategy to measure myelin with MRI.

To facilitate further progress, we provide a detailed characterisation of the evaluated studies as an online resource. We also share a set of 12 recommendations for future studies validating putative MR-based myelin markers and deploying them *in vivo* in humans.

**Highlights:**

- Systematic review and meta-analysis of studies validating microstructural imaging with myelin histology
- We find many MR markers are sensitive to myelin, including FA, RD, MP, MTR, Susceptibility, R1, but not MD
- Formal comparisons between MRI-based myelin markers are limited by methodological variability, inconsistent reporting and potential for publication bias
- Results emphasize the advantage of using multimodal imaging when testing hypotheses related to myelin in vivo in humans.

## 1. Introduction

Myelin is crucial for healthy brain function. Early studies found that myelin provides insulation and facilitates electrical conduction in neural circuits (Basser, 2004; Goldman and Albus, 1968; Waxman, 1980; Rush-ton, 1951). More recently, a host of observations have emphasised a wider set of roles for myelination, from enabling high frequency conduction (Saab et al., 2016) to providing trophic support for axons (Fünfschilling et al., 2012; Lee et al., 2012; Jensen and Yong, 2016). Changes in myelination occur as part of normal brain development (Gibson and Peterson, 1991; Ziegler et al., 2019) and aging (Peters, 2009; Hill et al., 2018). Moreover, recent studies have highlighted the possibility that myelination may change dynamically also in adulthood (Sampaio-Baptista et al., 2013), and that these changes may be crucial for learning and memory formation (Mckenzie et al., 2014; Pan et al., 2020; Steadman et al., 2020).

When myelin is damaged or lost (demyelination), neural communication is affected. Demyelination is a hallmark of many neurological diseases, such as multiple sclerosis, and affects white matter as well as grey matter (GM) (Lucchinetti et al., 2011). The possibility of inducing remyelination, either through pharmacological treatments (Stankoff et al., 2016) or behavioural interventions through experience-dependent myelin plasticity (Purger et al., 2016), is an emerging focus in clinical trials. It is also known that adverse events like social isolation can impact brain myelination (Liu et al., 2012; Makinodan et al., 2012) and myelination is involved in neuropsychiatric disorders including autism (Zikopoulos and Barbas, 2010) and schizophrenia (Stedehouder and Kushner, 2017). Therefore, robust *in vivo* markers to assess myelination could enable diagnosis and treatment monitoring for a wide range of conditions, as well as more detailed study of healthy brain function in humans.

MRI is the most frequently used tool to study brain structure and function *in vivo*, since it is noninvasive and widely available. The resolution of images acquired using a conventional MRI scanner lies in the millimetre range, whilst individual axons have diameters in the range of micrometres. Microstructural parameters can therefore only be estimated as summary measures and inferred from information obtained in comparatively large tissue volumes, e.g. cubic mm-sized voxels. This is possible because many aspects of MR physics are influenced by myelination (Does, 2018; Edwards et al., 2018; Möller et al., 2019; Novikov et al., 2019). Macromolecules in the myelin sheath influence relaxation rates (as measured by the longitudinal relaxation rate R1, the transverse relaxation rate R2, the effective transverse relaxation rate R2*, and myelin water fraction MWF) as well as magnetization transfer (MT; as quantified by the magnetization transfer ratio MTR, and the macromolecular pool size MP). The structure of the myelin sheath hinders local water diffusion (as measured by diffusion-weighted imaging (DWI)) and diamagnetic myelin influences the local magnetic field strength (as measured by quantitative susceptibility mapping (QSM)) (Möller et al., 2019; Weiskopf et al., 2015). In recent years a growing number of MR techniques have been successfully applied to study brain microstructure, and biophysical multi-compartment models have been developed to explicitly attempt to capture and quantify myelin-specific signals (MacKay et al., 2009; Mezer et al., 2013; Alonso-Ortiz et al., 2015; Campbell et al., 2017; Heath et al., 2018; Piredda et al., 2020).

Currently, it is not clear which of these MR markers is the most biologically accurate non-invasive measure for myelin, and how the measures differ in their sensitivity to specific aspects of myelination. While theoretical modelling can help in designing novel MR methods (Veraart et al., 2019), any model of MR signals will have intrinsic limitations given the complexity of brain tissue, including simplifying assumptions about the tissue geometry and the MR physics. For example, magnetization exchange between the modelled compartments is often assumed to be much slower than the MR measurements (Levesque and Pike, 2009; Barta et al., 2015; Does, 2018) and myelin water is often assumed to be the only driver of fast decay (Cohen-Adad, 2014). In light of these limitations, validation is a necessary step towards making MRI-based methods biologically interpretable, and applicable to clinical and basic research (Cohen-Adad, 2018; Barros et al., 2019).

Histological validation studies have been conducted for a variety of microstructural MRI metrics. Validity is often considered as the extent of agreement between a measured parameter and an underlying biological parameter of interest. For this reason, the gold standard for validation studies aiming to assess the accuracy of MR markers is to compare MRI and underlying myelin content within the same tissue. These studies perform MRI scanning in animals or in humans post-mortem, process the tissue histologically to obtain a ground-truth measure of myelination in the tissue, and then test whether variance in the microstructural MRI metric is driven by variance in myelination as assessed with histology. Neuroimaging studies conducted in vivo often use individual validation studies of this kind as evidence to justify employing a specific microstructural MRI measure to study myelination. However, validation studies use varying methodologies and have varying outcomes, making it difficult to assess to what extent such justifications are valid.

To better understand which MRI marker is best suited to measure myelin, we aim to collate evidence from the validation literature on microstructural MRI-metrics for myelin. First, we provide a comprehensive overview of validation studies reporting a correlation between MRI and histology. Second, we assess qualitatively a range of key methodological details known to influence histological signals (e.g. tissue processing), MR signals (e.g. state of the tissue during scanning) and correlation between the two (e.g. ROI definition method). Third, we perform meta-analyses to investigate how much variance is shared between each microstructural MR marker and histological myelin metrics. Fourth, we use insights from our systematic review to highlight the limitations of existing validation work and to develop a list of recommendations for future validation and in vivo imaging experiments.

## 2. Methods

### 2.1. Systematic review: study selection

A systematic review was conducted using PRISMA guidelines (Shamseer et al., 2015) and incorporating best practices from the AMSTAR 2 checklist for clinical meta-analyses where applicable (e.g. searching across multiple datasets, performing study selection in duplicate) (Shea et al., 2017). Articles on quantitative validation of MR markers for myelin were searched for in Pubmed and Scopus (search date: February 20, 2020), using the following search terms: (((myelin[Title/Abstract]) AND (post-mortem[Title/Abstract] OR post-mortem[Title/Abstract] OR histol*[Title/Abstract] OR *ex vivo*[Title/Abstract] OR histochem*[Title/Abstract] OR histopath*[Title/Abstract])) AND (MR*[Title/Abstract] OR magnetic resonance imaging[Title/Abstract])) AND (brain[Title/Abstract] OR spinal cord[Title/Abstract]). In addition, the literature was complemented with 45 articles from the authors literature library, which included references from recent reviews (e.g. Heath et al. (2018); Mancini et al. (2020)). These articles were subject to the same full-text screening procedure as the database-derived articles.

We included only peer-reviewed articles. We excluded review articles, phantom-only studies, atlas-based validation studies, MR spectroscopy studies, studies that do not quantify both myelin and MRI, and studies that do not perform quantitative comparisons of myelin and MRI. We did not specify exclusion criteria based on the investigated species or pathology or myelin validation method used.

All studies detected in the systematic search were screened for inclusion criteria in their abstracts. As most studies were excluded at this stage, abstract screening was independently run by two investigators (I.L. and A.L.) and discordant decisions on article inclusions were resolved by discussion and consensus-finding between the investigators. To verify the inclusion criteria based on the full-text of each article, a second stage of screening was also performed.

### 2.2. Systematic review: information extraction

To provide a comprehensive overview of the state of the validation literature for myelin, from each selected paper, we extracted information on the methodology of the validation aspect of the study. We chose methodological factors that may impact on the MRI-based metrics, the histological quantification of myelin or their correlation (also see Barros et al. (2019) on a similar review on iron imaging). The information is reported in Supplementary Tables 1 - 5 and is also openly available for external use at: https://lazaral.github.io/Myelin-Validation-Systematic-Review/. Below, we describe which information was selected and how it was reported.

**Table 1:**
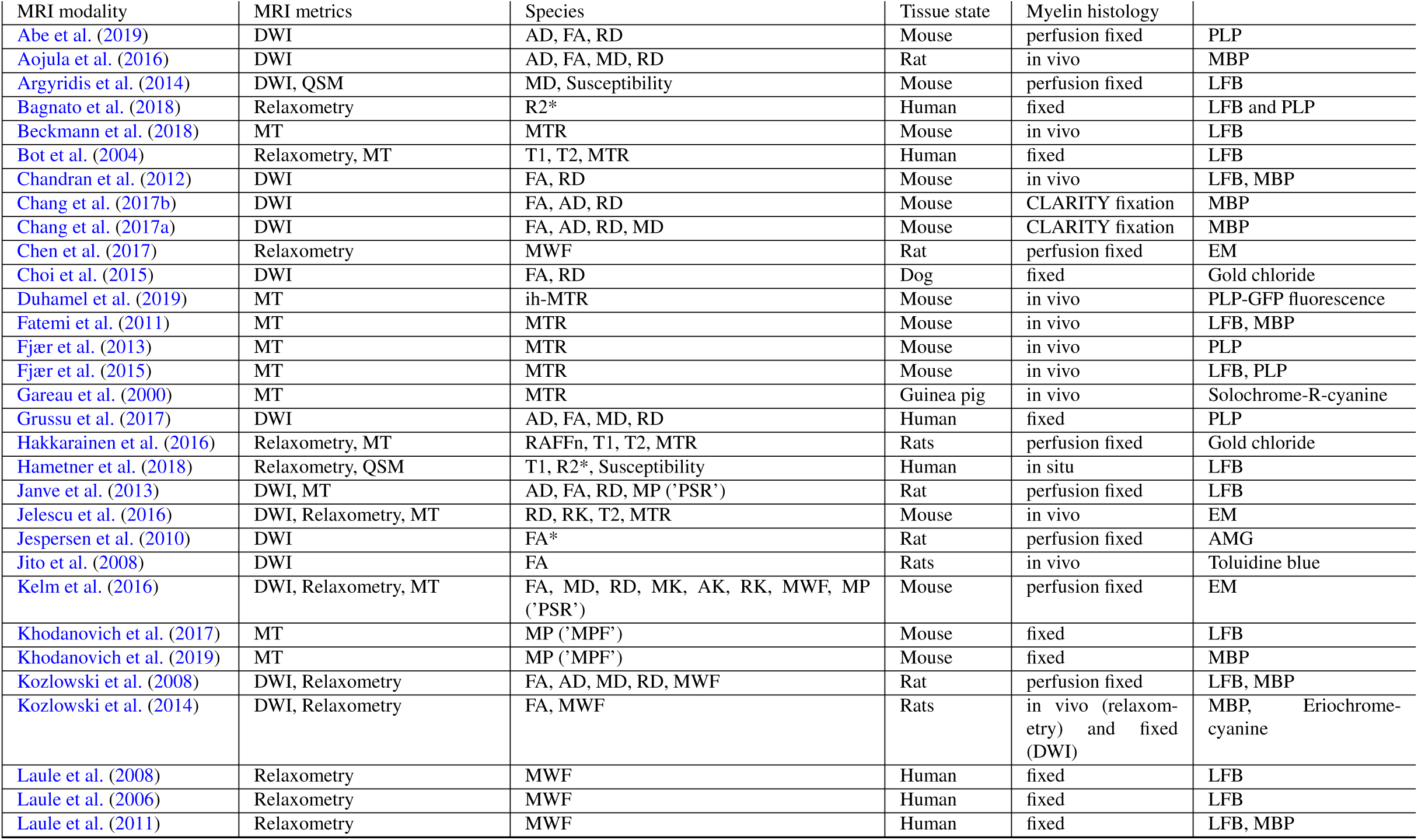
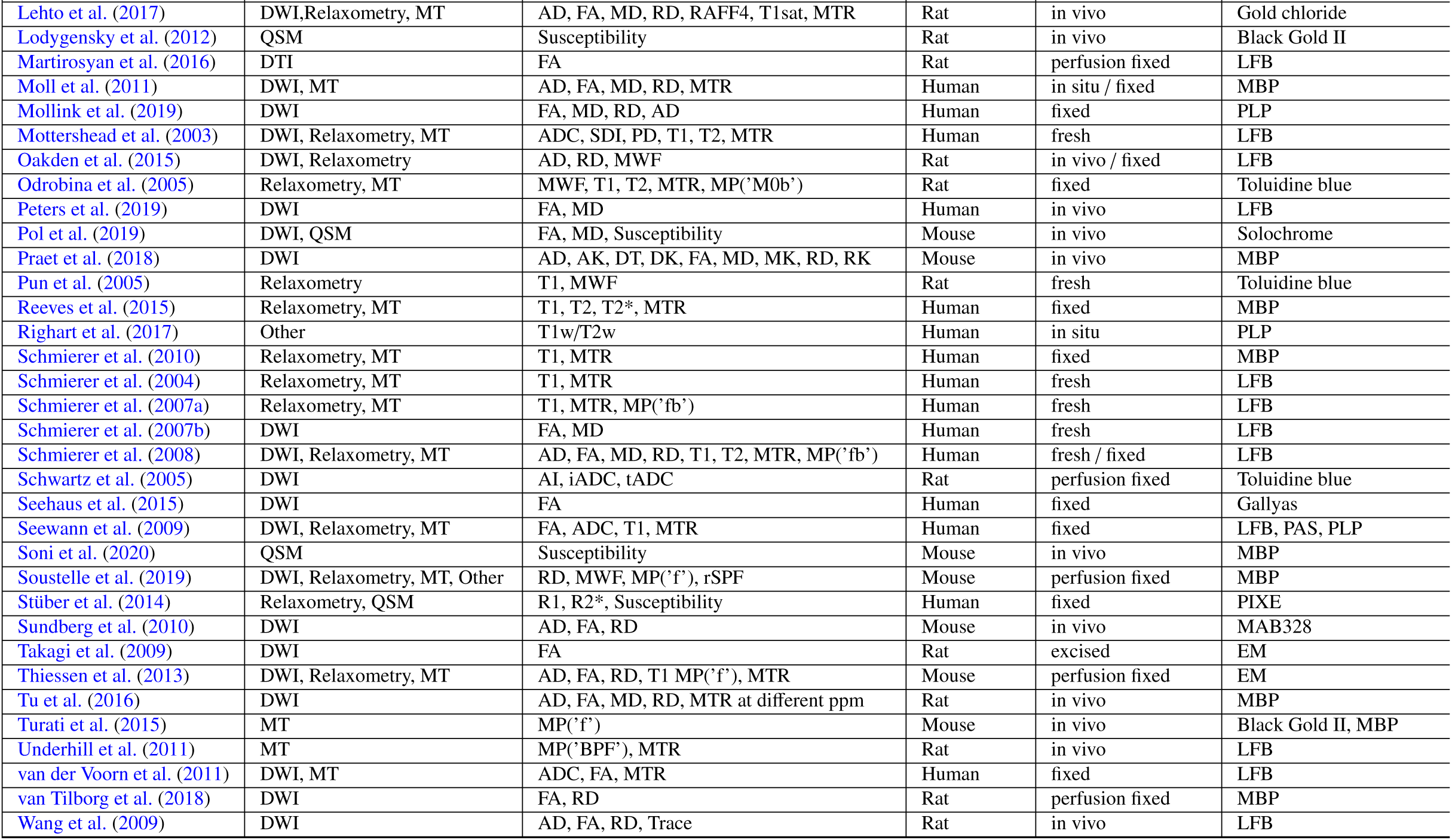
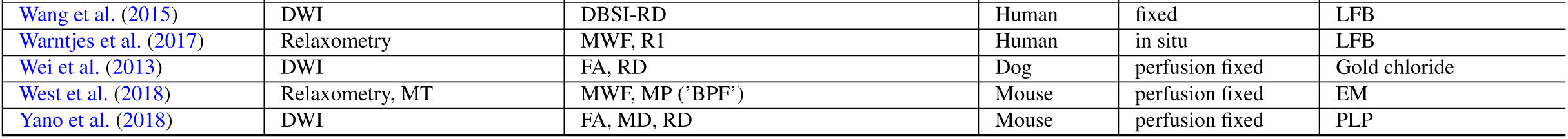
Overview of the assessed validation studies. Acronyms: DWI: Diffusion-weighted imaging: **AK:** axial kurtosis; **Al:** Anisotropy Index (tADC/lADC); **DBSI-RD:** diffusion basis spectrum imaging - based radial diffusivity; **DK:** diffusion kurtosis metrics; **DT:** diffusion tensor metrics; **FA:** fractional anisotropy (from diffusion tensor model); **1ADC:** longitudinal apparent diffusion coefficient (not modelled with tensor); **MD:** mean diffusivity (from diffusion tensor model); **MK:** mean kurtosis; **RD:** radial / transverse diffusivity (from diffusion tensor model); **RK:** radial kurtosis; **SDI:** diffusion standard deviation index; **tADC:** transverse apparent diffusion coefficient (not modelled with tensor); **Relaxometry: MWF:** myelin water fraction; **Rl:** longitudinal relaxation rate; **R2*:** effective transverse relaxation rate; **RAFF4:** Relaxation Along a Fictitious Field in the rotating frame of rank 4; **Tl:** longitudinal relaxation time; **T2:** transverse relaxation time; **T2*:** effective transverse relaxation time; **MT: magnetisation transfer: BPF:** bound pool fraction; **F:** pool size ratio; **Fb:** macromolecular proton fraction; **ih-MTR:** MTR from inhomogeneous MT; **MOb:** fraction of magnetization that resides in the semi-solid pool and undergoes MT exchange; **MP:** macromolecular pool; **MPF:** macromolecular proton fraction; **MTR:** magnetisation transfer ratio; **PSR:** Macromolecular-to-free-water pool-size-ratio; **STE-MT:** MTR based on short echo time imaging; Tlsat: Tl of saturated pool; **UTE-MTR:** MTR based on ultrashort echo time imaging; **QSM:** quantitative susceptibility mapping; **Others: rSPF:** relative semi-solid proton fraction from an 3D ultrashort echo time (UTE) sequence within an appropriate water suppression condition; **Tlw/T2w:** ratio of image intensity in a Tl-weighted vs T2-weighted acquisition.AMG: Autometallographic myelin stain; **LFB:** Luxol fast blue stain; MA: fraction of myelinated axons; **MAB238:** Anti-oligodendrocyte immunohistochemistry; MBP: Anti-myelin-basic-protein immunohistochemistry; **PAS:** periodic acid-Schiff; **PIXE:** proton-induced X-ray emission; **PLP:** Anti-proteolipid-protein immunohistochemistry.

#### 2.2.1. Imaging: modality, parameters and metrics

Studies used one of five imaging modalities: a) **Relaxometry**: acquisitions aimed at estimating relaxation times / relaxation rates or myelin water fraction based on its short relaxation rate; b) **DWI**: metrics based on diffusion models (these metrics are known to be influenced by both myelin and non-myelin factors, but are often used as microstructural markers (Pierpaoli et al., 1996; Beaulieu, 2014; De Santis et al., 2014) c) **MTI**: acquisitions applying off-resonance pulses to induce magnetization transfer effects and aiming to quantify them; d) **QSM**: metrics based on the quantification of susceptibility based imaging; and e) **Others**: acquisition that do not fall into categories a-d, such as the T1w/T2w ratio (Glasser and Van Essen, 2011).

The quantitative MT models that aim to assess the proportion of macromolecules (assumed to be mostly macro-molecules in myelin) in a voxel all use different terms for this parameter. While f denotes the ratio of the macro-molecular water pool over the free water pool, the fraction (ratio of the macromolecular water pool divided by the sum of all pools) is denoted in a number of ways. We summarize this metric as macromolecular pool (MP) and indicate in brackets the original term used in the table.

In addition to the metrics quantified, we considered the field strength at which the imaging was done and the resolution of the acquisition protocol. If voxel resolution was not reported, it was deduced from FOV and matrix size. We report all resolutions in mm for comparability across studies, rounded to three significant figures.

#### 2.2.2. Tissue types

We extracted information for what species was used, which part of the central nervous system was imaged (e.g. brain vs spinal cord, whole brain vs sections or smaller samples) and whether it came from healthy individuals or natural or induced pathologies.

We additionally extracted information on what anatomical structures or tissue types were used for the statistical analysis specifically. Depending on the level of detail provided by the paper, this was either anatomical regions of interest (ROI) or tissue types (e.g. identified by degree of pathology). We also extracted whether manual or automatic delineation of ROIs was performed. We classified the ROI definition as manual if words like outlined or drawn or defined or defined and labeled or designated were used without further mentioning of an automated or semi-automated tool. If atlases were used to segment ROIs, the specific atlas used is reported. If voxelwise analysis statistics were computed, the ROI definition was set to NA (not applicable).

#### 2.2.3. Tissue preparation

As stage and type of tissue fixation both affect MR parameters (Dusek et al., 2019), we extracted information on the state of the tissue during MR imaging. We distinguished between in vivo, in situ, fresh, fixed. If scanning was not done *in vivo*, we also looked at post-mortem times (defined as time from death to fixation), which are indicative of the autolytic state of the tissue at the time of fixation. Post-mortem time was reported as NA (not applicable) when scanning was done *in vivo* and/or in perfusion fixed tissue. If it is reported with±sign, then this indicates mean and standard deviation across samples.

As temperature has an effect on MR relaxation times (Birkl et al., 2016) and diffusion (Dhital et al., 2016), tissue temperature during scanning was extracted. If scanning was performed *in vivo*, body temperature was assumed. Last but not least, we report how tissue was treated for histology (cryosectioning or paraffin embedding), which can affect tissue staining success and intensity (Werner et al., 2000).

#### 2.2.4. Histology methods

Given the diversity of potential histology methods, we extracted information on how myelin was histologically visualised, including staining methods.

Some microstructural features of the brain, such as iron and axonal density, are related to myelin and have similar effects as myelin on MR signals (Möller et al., 2019). We assessed whether iron or axons have also been considered by the studies we reviewed, and report this information in the form of binary columns in the table). This was determined by searching the text for any mention of histological methods that could be used to estimate iron or axons (e.g. iron stains, anti-neurofilament immunohistochemistry or electron microscopy).

We extracted the histology quantification method, which belonged to one of the following categories: a) **Staining fraction**: this category was chosen if the microscopy image was segmented and the fraction / percentage of area stained was quantified, b) **Staining intensity**: this category was chosen if the optical density of one or more colour channels was quantified, c) **Inverse staining intensity**: similar to staining intensity, but instead light transmittance was quantified, leading to a different expected direction of the correlation coefficient between MRI and histology.

We also report the thickness of the sections used for histology (all reported in *μ*m,converted if necessary for the sake of comparison across studies).

#### 2.2.5. Statistics

An inclusion criterion for our screening was that quantitative correlation between MRI and histology must have been performed. Therefore, we extracted the specific type of correlation that was used in each paper - whether average values within a given ROIs were compared between histology and imaging, or whether pixel-wise or voxel-wise correlations were employed.

We then considered what design was used for the main correlation analysis. Validation studies can either focus on whether for the same subject differences in MR metrics across ROIs are reflected in histological metrics (within-subject validation design, also known as spatial correlation) or whether for the same ROI, differences in MR metrics between subjects are reflected in histological metrics (between-subject validation design). Therefore, we distinguished between a) **Between-subject**: correlations where each data point originates from a separate sample/subject; b) **Within-subject**: correlations where all data points originate from the same sample/subject (see Supplementary Figure 1 for a schematic illustration of the difference between between-subject and within-subject design); c) **Mixed (modelled)** - designs where all ROIs across all subjects were included in the same analysis, taking into account that multiple data points were derived from the same subject. By formally correcting for the between-subject or within-subject variance in the data, these studies effectively report either within-subject or between-subject correlation coefficients. d) **Mixed (not modelled)** - all ROIs across all subjects included in the same analysis, without taking into account that multiple data points were derived from the same subject. In these studies, it is impossible to disentangle whether the correlation coefficient is driven by within- or between-subject variance.

**Figure 1:**
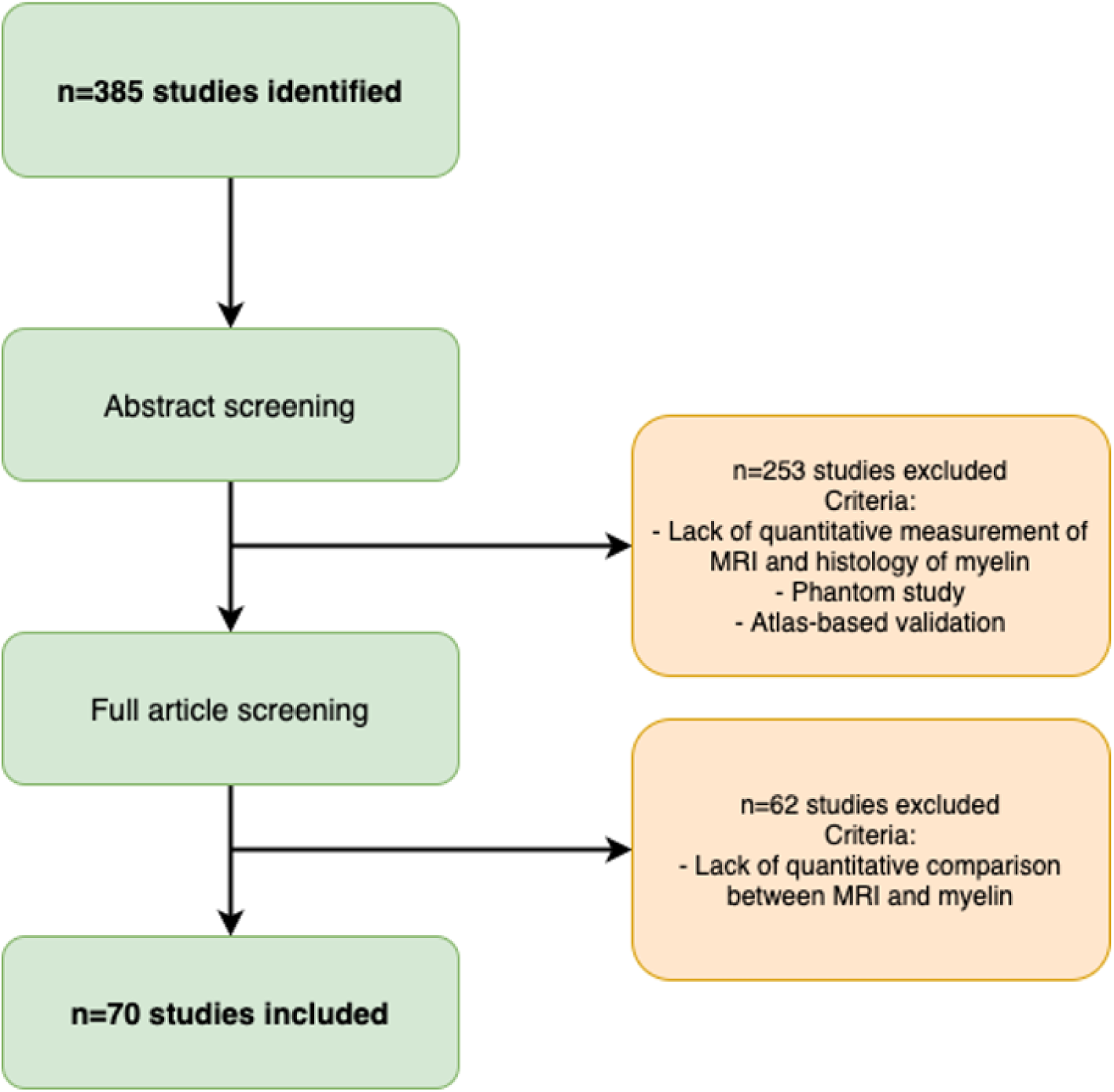
Flow diagram of study selection.

We extracted the sample size used for the correlation analysis. We reported sample size per group (number of subjects in each group, e.g. in control and in disease group), as well as total subjects (total subjects across groups). For studies using multiple ROIs, the effective sample size for the correlation differed from the number of subjects used. Therefore, we also reported the number of ROIs used. In Within-subject studies with varying numbers of ROIs for each individual slice/sample, the range of ROIs for each slice/sample was reported. In Mixed (unmodelled) statistical approaches, often only the total number of ROIs across all subjects is reported in the article, and we reported this figure instead.

The matching of histology and MRI was also considered. No co-registration indicates that ROIs were specified in native space in histology and MRI. NA (not applicable) was used if registration is not feasible, such as for electron microscopy, where the field of view for microscopy is generally only a small fraction of the field of view for MRI. If MRI and histology data were coregistered, we extracted the type of registration (e.g. manual, affine transform, etc.)

For the statistical results, we reported statistical methodology (Pearson correlation, Spearman correlation or linear regression), correlation effect size, significance of the correlation, and regression slope (if applicable, otherwise NA). Statistical methodologies including mixed effect models and multilevel models were all labelled as linear regression. Reported effect sizes included Pearson or Spearman correlation coefficient (directional) or a coefficient of determination *R*^2^ (non-directional). If regression was run but no linear equation details were reported, we indicated this with not reported in the table.

### 2.3. Meta-analysis: study selection

Meta-analyses aim to estimate effect sizes across multiple studies. However, only comparable metrics and statistical designs can be pooled together. For example, different meta-analyses need to be run for papers reporting Spearman and Pearson correlation coefficients. Moreover, studies using within-subject and between-subject designs need to be considered in different analyses. Therefore, we excluded from our meta-analysis studies where correlation coefficient type could not be determined, and studies using a mixed design where either within- and between-subject variance could have determined the outcome.

Within included studies, separate meta-analyses were carried out for any MRI marker that was validated in more than two independent studies with more than three subjects. For studies with between-subject designs using separate ROIs or separate groups of subjects, the correlation coefficients were averaged (using Fishers r-to-z transformation) before entering the meta-analysis. If correlations with separate histological markers were reported as results, the results were fed separately into the metaanalysis.

### 2.4. Meta-analyses: statistical analysis

All meta-analyses were run through the *meta* R package (Schwarzer et al., 2007). The coefficient of determination *R*^2^ was used as a common effect size measure and a fixed-effect model was employed to weigh different studies based on their sample size (Schulze, 2005). For each MR modality, a forest plot was used to summarise studies included, study sample sizes, and effect sizes of individual studies as well as meta-analytic effect size (with 95% confidence intervals). A confidence interval that had a range of exclusively positive values was considered indicative of evidence for a correlation between MRI and histology.

### 2.5. Meta-analyses: p-value distribution in the literature

Questionable Research Practices, whether performed consciously or unconsciously, can lead to skewed effect sizes in the literature. For instance, as significant studies are more likely to be published, there can be both conscious and unconscious biases leading researchers to report biased *p*-values (Button et al., 2013). The effects of this pressure to find significance are reflected in the literature: although *p*-values would be expected to appear randomly in the literature, p-values right under 0.05 are over-represented in the psychology literature (Head et al., 2015). To test whether the validation literature in our meta-analyses is biased, we perform a *p*-curve analysis through the *dmetar* R package (Simonsohn et al., 2014; Harrer et al., 2019).

### 2.6. Meta-analyses: quantifying and correcting for publication bias in the literature

Funnel plots are a common tool to estimate publication bias in meta-analyses (Egger et al., 1997). For any given effect, a meta-analysis calculates, one would expect high-sample-size studies to best approximate that effect (visualised by the peak of the funnel). In contrast, low-sample-size studies are expected to provide noisier estimates of the true underlying effect size, leading to more variance in the effect size estimations and thus forming the wide part of the funnel as effect sizes decrease.

While funnel plots can capture the presence of publication bias, they are also sensitive to other effects. For example, an asymmetric funnel plot can arise from poor methodology in small-sample-size studies, heterogeneous true effect sizes across studies, or appear by chance (Sterne et al., 2011). However, funnel plots have the advantage that by quantifying potential publication bias, they allow correcting for it. To demonstrate the robustness of our results to potential publication bias, we perform trim-and-fill procedures based on (Duval and Tweedie, 2000) and implemented through the *trimfill* function in the *dmetar* R package.

Finally, we also complement funnel plots with Spearman correlations between sample sizes and effect sizes across the studies included in the meta-analysis.

### 2.7. Data and code availability

All extracted information and quantitative meta-analysis code is openly available at https://lazaral.github.io/Myelin-Validation-Systematic-Review/.

## 3. Results

### 3.1. Exploring the features of the MR-histology validation literature

Our search yielded 385 unique articles (Figure 1). Of these articles, 294 were from Pubmed, 46 from Scopus, and 45 from our expert library. Abstract screening led to exclusion of 2532 studies, while full-text screening further excluded 62 studies, with a total of 70 remaining studies fitting our inclusion criteria (Table 1). We believe this constitutes the state of the field at the time of writing.

Information from the included studies was extracted and summarised in Supplementary Tables 1, 2, 3, 4, 5. To have a better overview of the literature, we provide quantitative summaries of our findings on the literature content (Figure 2). We find that validation studies have a median sample size of 13 (Figure 2A), comparable to the median of the most cited fMRI studies during the period of publication included in our meta-analysis (12), but below current median sample size (20) (Szucs and Ioannidis, 2020). The validation literature uses a wide range of histological markers (with 9 studies using more than one marker, Figure 2F). It also employs a wide variety of species and field strengths used for experiments (Figure 2D). Taken together, these factors highlight that the validation literature has considered different field strengths, species, and MRI and histology approaches to quantify myelin, making common results across papers highly generalizable.

**Figure 2:**
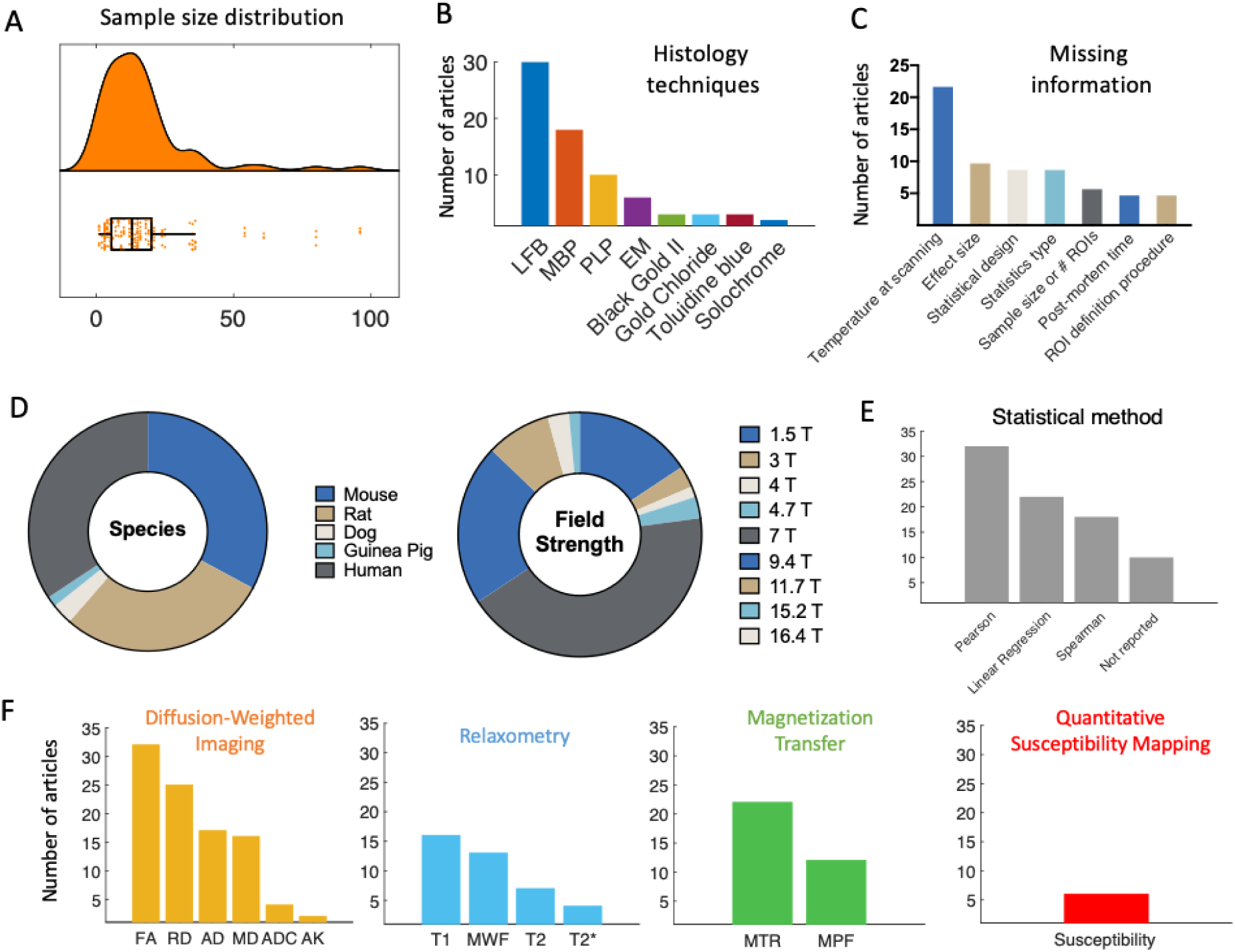
Features of the MR-histology validation literature. **[A]** Sample size distribution of the included papers (median = 13 subjects). **[B]** The most frequently used histology techniques used in the articles are shown in decreasing order. **[C]** Due to lack of commonly agreed reporting standards, some relevant information may be missing from validation studies. The most commonly missing pieces of information on methods are reported here. **[D]** Pie chart of species and field strength used in each study. **[E]** Frequency of different statistical measures used for the validation analysis. **[F]** MR markers most commonly used in the literature, subdivided based on MR technique they are based on. The graphs show that DWI, MT and relaxometry are by far the most commonly explored techniques in validation studies. Within studies using DWI, tensor-based measures are predominant.

The studies we included test the validity of a wide-range of potential myelin markers, reflecting the range of markers used in application studies. For instance, the most commonly validated MR markers are DWI-based (such as FA), with relaxometry and MT-based studies close seconds (Figure 2F). We also note that many markers appear only once in the literature (e.g. RAFF4, ih-MT), suggesting there are many more markers in the literature that lack thorough validation. We do not include these markers in Figure 2F for simplicity.

Finally, we observed that many studies fail to report some crucial aspect of their methodologies or results. We therefore reported the most commonly missing experimental details (Figure 2C).

### 3.2. Meta-analysis

A total of 32 studies were selected for inclusion in a quantitative meta-analysis based on this selection process (24 reporting Pearson only, 7 reporting Spearman only, and 1 reporting both; 22 using between-subject design, 10 using within-subject design). To ensure comparability between studies included, separate meta-analyses were run for studies using between-subject and within-subject variance (Figure 3 and Supplementary 5 respectively). Moreover, studies using Pearson and Spearman coefficients were also pooled separately (meta-analyses are labelled Pearson or Spearman within Fig.3). On the included studies, we ran fixed-effects meta-analyses to estimate a cross-study validation effect size for each marker (Figure 3). Except for mean diffusivity, we find meta-analytic evidence for correlations between histological markers and all markers investigated, with no clear marker having a stronger effect size than others.

**Figure 3:**
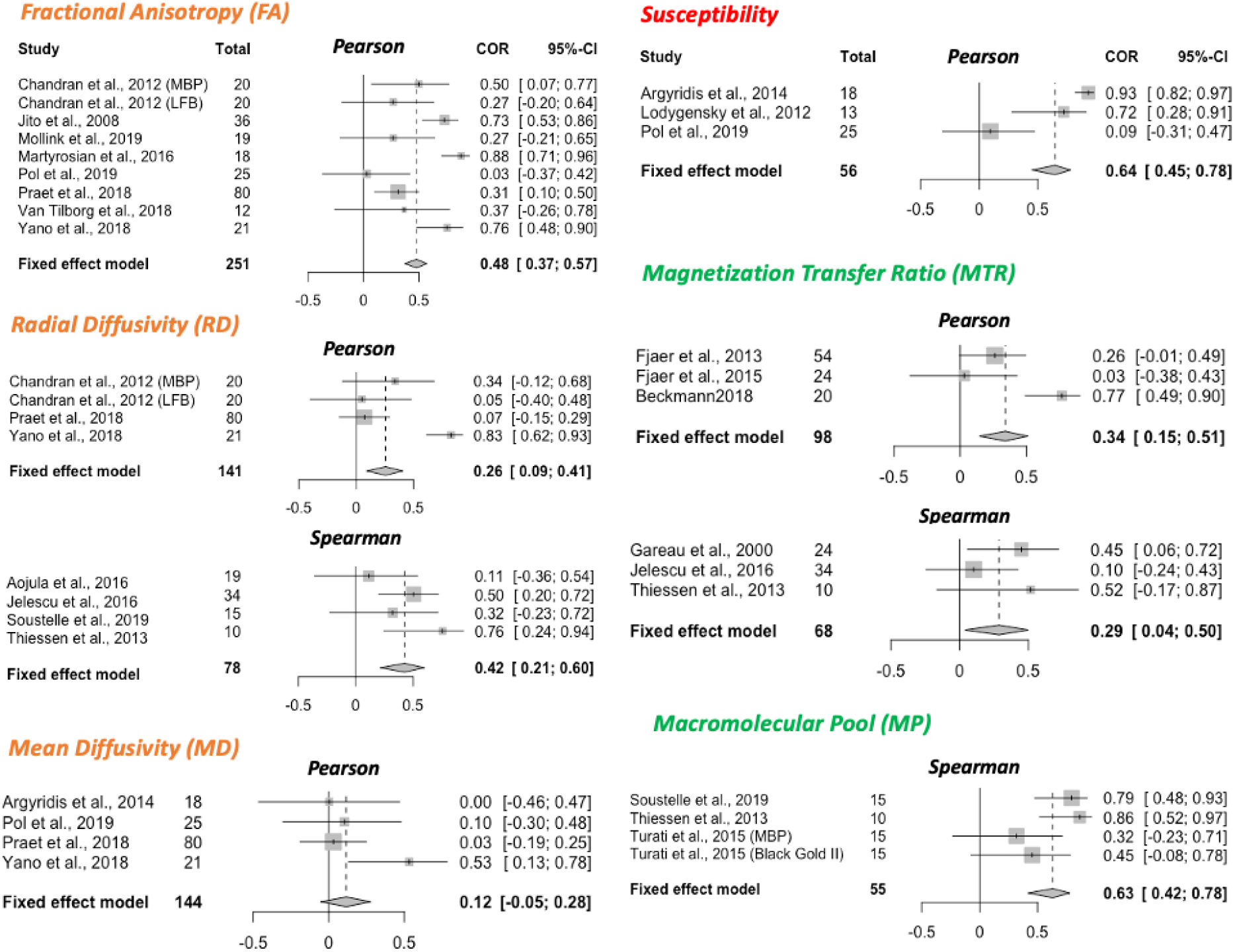
Meta-analysis of correlations between histology and MR markers. Forest plots from meta-analyses of 6 MR markers (FA, RD, MD, susceptibility, MTR and MP). Spearman and Pearson correlations are pooled separately; the type of correlation coefficient is indicated at the top of each forest plot. All studies included use a between-subject statistical design; meta-analyses of studies with a within-subject design are reported in Supplementary Figure 5. For each study, sample size and effect size (*R*^2^) with the respective confidence interval are provided. The fixed effect model results are reported at the bottom of each forest plot, together with pooled sample size across all included studies and with 95% Confidence Interval of the meta-analytic effect size.

### 3.3. P-value distribution bias and Publication Bias

Within the studies included in our meta-analyses, we then test for sources of bias. We find no evidence of abnormal p-value distribution, with a left-skewed p-curve and 80% of positive results providing p-values of 0.01 or lower (Supplementary Figure 2). This indicates that conscious and unconscious bias towards significant results has likely not affected the study outcomes in the literature.

We then test for evidence of broader publication bias through a funnel plot (Supplementary Figure 3). We find a significant correlation between sample size and effect size, where higher effect sizes in the literature are reported from studies with the smaller sample sizes.

While asymmetric funnel plots can arise from true heterogeneity in effect sizes across studies (Sterne et al., 2011), which may be the case in our studies, we perform a sensitivity analysis to test whether correcting for such asymmetry would alter the results. We perform trim-and-fill procedures (Supplementary Figure 4) based on (Duval and Tweedie, 2000) and find that if present, publication bias would inflate the validation effect size of some markers.

## 4. Discussion

#### Recommendations for in vivo human studies aiming to measure myelin with MRI and for future validation studies

##### 1. Recommendations for in vivo human studies aiming to measure myelin with MRI

- 1.1 Use acquisition protocols with multiple myelin-sensitive modalities (multimodal imaging)
- 1.2 Balance evidence from theoretical MR modelling and from histological validation studies when selecting MR markers to test hypotheses on myelin
- 1.3 Take validation parameters into account when selecting an MR marker for your study: a marker validated only in within-subject studies may not be sensitive to between-subject variability in myelination, and vice versa

##### 2. Recommendations for experimental protocols in validation studies

- 2.1 Report fixation protocols, and ideally use established ones
- 2.2 Monitor and report scanning temperature
- 2.3 In human studies, report and account for post-mortem time
- 2.4 Use myelin histology specific for the experiment’s objective, and where needed probe histologys specificity to the phenomenon of interest
- 2.5 Use automated ROI definitions and co-registration between MR and histology

##### 3. Recommendations for statistical analyses in validation studies

- 3.1 Take the subject structure of the data into account ROIs from the same subject are not independent, and multiple ROIs from multiple subjects cannot be pool together without modelling the nested structure of the data
- 3.2 Use correlation methods robust to outliers, and check distributional assumptions
- 3.3 Covary for non-myelin factors such as axon density and iron
- 3.4 Pre-register analyses and share data to reduce the impact of publication bias

Our systematic review yielded 70 validation studies of microstructural MRI metrics to measure myelin, which included a wide range of MR markers and histological techniques. We then performed a detailed qualitative characterisation of this literature, as well as a quantitative meta-analysis of the reported effects sizes.

### 4.1. Many MRI markers correlate with myelin

Our meta-analyses of 7 commonly used MR markers (FA, RD, MD, susceptibility, R1, MTR and MP) show that there is evidence all these markers correlate with myelin, with the exception of MD. The effect sizes for each of these markers are pooled across a heterogeneous literature, which highlights the robustness of MR myelin markers to features of individual validation studies, such as field strength, animal models, and histological measure. This indicates that correlations between histological and MR-based markers of myelin are not restricted to any one setting, and is promising for translational uses of MR-based markers in contexts where obtaining histology is not possible.

### 4.2. Challenges in measuring myelin for in vivo studies in humans

One important question that the validation literature aims to tackle is whether one MR marker outperforms others in how well it captures myelin signals from a tissue. While some markers show stronger meta-analytic effect sizes than others, it is difficult to infer whether different markers relate to myelin to different extents. In particular, two key issues prevent making conclusive comparisons on how different MR markers compare with each other. One is the difference in methodologies between studies - effect sizes may be influenced by factors ranging from tissue processing to scanning temperature, and the small number of studies available makes it difficult to systematically account for these factors. The other is the inconsistent reporting: studies report results inconsistently (e.g. in terms of reported statistics) and sometimes fail to report crucial details.

In this respect, our results underscore the importance of employing multimodal acquisitions, as observing similar effects across multiple markers is at present the best way to verify hypotheses related to myelin **(Recommendation 1**.**1)**. Techniques to explicitly combine multiple imaging modalities already exist (e.g. joint inference (Winkler et al., 2016), multimodal combination Mangeat et al. (2015)), and many studies have already found that multiple myelin markers can provide complementary information (Lipp et al., 2019; Eichert et al., 2020).

Another crucial aspect highlighted by our results is the need to balance evidence from theoretical models with evidence from validation studies, when selecting MR markers for a human in vivo study on myelination **(Recommendation 1**.**2)**. This is particularly clear for diffusion MRI, where advanced models, such as the Spherical Mean Technique (Kaden et al., 2016) and neurite orientation dispersion and density imaging (NODDI; Zhang et al. (2012), are becoming increasingly popular compared to tensor-based metrics such as FA, but are vastly under-represented in the validation literature (Alexander et al., 2019; Caspers and Axer, 2019). Therefore, it will be key to the continued success of these new methods to verify what they are sensitive to at the cellular level. In particular, well conducted validation studies will be critical to confirm the improved microstructural sensitivity of these markers compared to markers from simpler tensor models.

A notable absence from our meta-analyses is Myelin Water Fraction (MWF). While 13 studies in total examine MWF, a high number of these studies use a mixed within- and between-subject design, or do not report the type of static measure used, which makes them difficult to aggregate in a meta-analysis. Therefore, further validation studies for MWF, and especially studies examining between-subject variance, may be needed.

A final recommendation for human in vivo studies is to take validation parameters into account when selecting which MR marker is most appropriate for a given study **(Recommendation 1**.**3)**. Some markers are only validated with one type of statistical design, which limits their potential applications. Relaxometry metrics such as R1 are generally more commonly validated with within-subject designs, which means it is unclear whether they would be able to pick up between-subject variability in myelination in the general population. Therefore, microstructural studies computing brain-behaviour correlations, where between-subject variability in myelination is key to the hypothesis tested (Johansen-Berg, 2010), might benefit from including markers that have been validated in between-subject designs in order to maximise their sensitivity to the phenomenon of interest.

### 4.3. Challenges in acquiring MRI data for validation studies

MR signals are affected by a variety of data acquisition choices. For instance, we find that the studies in our review validate MR metrics with either *in vivo* or *ex vivo* MR scanning, each of which has advantages and disadvantages. For studies using human brains, the option to combine *in vivo* scanning with *ex vivo* histology is rare (see Treit et al. (2019)), whereas in animal studies, both *in vivo* and *ex vivo* scanning are possible. *In vivo* scanning has the advantage that it better resembles the parameters of *in vivo* human studies, while *ex vivo* scanning has the advantage that the tissue is scanned and processed for histology in a similar state. Also, higher resolution can be achieved with *ex vivo* imaging, which in studies of animals with smaller brains provides more anatomically comparable detail to human studies with larger voxel sizes (Lerch et al., 2012).

#### 4.3.1. Ex vivo tissue fixation affects microstructural MRI

Tissue fixation is a necessary step for stopping tissue decomposition in *ex vivo* scans, but it also has a major influence on MR signals. Formalin fixation and the embedding medium for scanning can affect image quality and MRI parameters (Dusek et al., 2019), even when excess fixative is being washed out of the tissue before scanning. This is because fixation works by linking amino acids in proteins, which means that the molecular tissue structure is inherently changed during the process (Thavarajah et al., 2012). Moreover, MR relaxation relies on water protons behaving differently depending on their molecular environment, and fixation does change this environment, for example by affecting membrane permeability (Shepherd et al., 2009).

Fixation-dependent changes in MRI have two effects on validation studies. First, if validation evidence only comes from fixed tissue, the generalisability to fresh tissue is not guaranteed. Among the studies we analysed, only a few of them assessed tissue that was scanned fresh, whereas ideally, both states should be validated (e.g. Schmierer et al. (2008)). Second, the variability in the type and duration of fixation can lead to variability in the MRI-based metrics. Longitudinal studies investigating tissue fixation report spatially and temporally varying effects on relaxation rates, MWF and also MP (Dawe et al., 2009; Seifert, 2019; Shatl et al., 2018). In the majority of studies we assessed, the fixation process ranged from a few days to a few weeks, which means the MR signal was likely very different between studies. Within an individual validation study, this influence may be minimized by using a standardised tissue processing protocol for all samples **(Recommendation 2**.**1)**. However, systematic differences of fixation effects across different ROIs may still pose problems for the quantitative comparison between MRI and histology.

#### 4.3.2. Scanning temperature affects microstructural MRI

Another factor to consider is temperature, which affects MRI parameters (e.g. Birkl et al. (2016); Dhital et al. (2016)). When scanning in vivo, the tissue is naturally kept at body temperature. However, the temperature conditions during post-mortem scanning are more flexible, allowing to either scan at room temperature or at a controlled temperature. Importantly, the tissue temperature could unintentionally change throughout the scanning session due to gradient heating, making it important to monitor scanning temperature in *ex vivo* scans. If the temperature varied across samples or across ROIs within the same sample, this could induce unwanted variability in the MRI metrics. In the assessed *ex vivo* validation studies, the scanning temperature was frequently not reported, making it difficult to estimate its impact on the results and to compare microstructural MRI metrics across studies **(Recommendation 2**.**2)**.

### 4.4. Challenges in obtaining ground-truth histological data for validation studies

#### 4.4.1. No standard pipeline for histological quantification exists

Validation studies aim to obtain a ground-truth measure for myelin content, but we find that pipelines used to obtain myelin content histologically varied widely across the assessed papers. The vast majority of microscopic visualisation employed classical stainings (most frequently LFB and Gold-based stains) or immunohistochemistry approaches (most frequently anti-MBP and anti-PLP). These approaches have primarily been developed to visualise myelination patterns in the tissue, rather than quantify myelin content (e.g. see Carriel et al. (2017a); Kiernan (2007); Thetiot et al. (2018); Woodhoo (2018)). Their advantage is that they allow imaging large fields of view, which is often needed in a validation study to compensate for the comparatively low spatial resolution of MRI relative to microscopy.

To analyse the resulting microscopy images, the majority of the assessed validation studies used either staining fraction (the relative amount of stained tissue) or average staining intensity in an area (the total amount of stain in the tissue) for quantification. However, both metrics come with limitations. Staining intensity is often assumed to scale linearly with myelin content. However, this is not always true, for example in cases where staining saturation can take place (e.g. in silver staining of WM (Pistorio et al., 2006)). Moreover, it is known that different samples (or different anatomical areas within the same sample) are not affected homogeneously by preparation steps such as fixation or antibody penetration (Dawe et al., 2009; Seehaus et al., 2015). Using staining fraction to quantify myelin could circumvent this issue, if the image segmentation intensity threshold is adjusted flexibly (Mollink et al., 2019), depending on local intensity variations. However, it remains poorly understood to what extent staining fraction values are comparable across different staining methods.

One solution to address the limitations of these commonly used metrics would be to employ methods developed for quantification, rather than visualisation, of molecules. These methods include proton induced X-ray emission (PIXE, Stüber et al. (2014)) and lipid mass spectrometry (Gonzaález de San Romaán et al., 2018), which could provide quantification of myelin for each imaging pixel, for example by estimating myelin concentration from sulfur and phosphorus (Stuüber et al., 2014). With the exception of one study (Stüber et al., 2014), these methods have not been used in the validation papers we reviewed, but may be used in future studies to provide complementary information to traditional histological techniques.

Another disadvantage of traditional histology methods is that what they gain in field-of-view size they lose in resolution. Higher-resolution histological methods such as electron microscopy and coherent anti-Stokes Raman scattering microscopy can visualise even individual myelin sheaths within small tissue blocks (generally not more than a few mm of size). With sufficient image quality and appropriate image segmentation (Stikov et al., 2015b; Zaimi et al., 2016), these methods have the potential to estimate the myelin fraction of imaged tissue, as well as the packing density and total surface area of the myelin sheaths. While they are not as common in the validation literature, and often not feasible over large areas of the brain, these methods provide more detailed information compared to stainings or immunohistochemistry, and could provide further validation for microstructural MR markers (Sternberger et al., 1978; Vincze et al., 2008).

#### 4.4.2. Tissue conditions influence histological quantification

The success of any histological method does not only depend on its molecular principles underlying myelin visualisation, but also on experimental factors, such as tissue processing protocols (e.g. Werner et al. (2000)). In our review, processing of the tissue for histology varied considerably across studies, with both paraffin embedding and cryosectioning being used with about equal frequency. While tissue fixation or freezing do not have a strong effect on lipids (Carriel et al., 2017a), extraction by solvents, such as used in paraffin embedding, extracts most lipids and only retains those covalently bound to protein (Kiernan, 2007; Carriel et al., 2017b). The myelin structure in the stained tissue is therefore inherently affected by how it was processed for histology and may not be comparable across studies that used different processing strategies **(Recommendation 2**.**1)**.

In studies with human tissue, a further complication is that post-mortem time also influences histological staining. The length of the post-mortem time indicates the advance of the tissue autolysis and therefore the microstructural intactness of the tissue (Sele et al., 2019), but it can vary between a few hours and a few days between samples. While a considerable number of assessed papers did not report post-mortem times, in those who did, it varied between 4 hours and 3 days. Large variability in postmortem times across tissue samples may induce unwanted variability in the histological quantification, and may need to be taken into account during the analysis as a covariate **(Recommendation 2**.**3)**.

#### 4.4.3. Histological specificity for myelin may depend on the experimental context

In our systematic review, most of the validation studies assessed only used one histological method. However, as different histological methods rely on wide-ranging and sometimes poorly understood molecular mechanisms, the agreement between MRI and histology will also depend on which histological marker is used. In the few papers that used multiple histological methods, validation results were not always the same for the different histological markers used (e.g. Kozlowski et al. (2008); Laule et al. (2011)). This may be because the extent to which the histological quantification is an accurate representation of myelin content might differ based on the characteristics of the sample or pathology in question. Myelin is a highly complex substrate, with the dry mass consisting of about 70% lipids and 30% proteins (Gopalakrishnan et al., 2013; O’Brien and Sampson, 1965). Its composition can vary between tissue types (Gonzaález de San Romaán et al., 2018), species (Gopalakrishnan et al., 2013) and in pathologies (Wheeler et al., 2008), and therefore different visualization strategies may be useful for different scenarios.

For example, immunohistochemistry can be used to target specific myelin constituents such as MBP and PLP, using antibodies that are attached to (most often through secondary antibodies) visualising elements, such as fluorescent molecules or diaminobenzidine tetrahydrochloride. If the target molecules are affected in the specific sample studied, this will affect the visualisation results. In studies using Shiverer mice, which have a genetic mutation for the MBP-gene (Molineaux et al., 1986), MBP-immunohistochemistry is well suited to demonstrate the genetic intervention, but could lead to a biased myelin quantification when used on Shiverer mice in a validation study.

Unlike immunohistochemistry, classical myelin stains are not specific to individual macromolecules of myelin, but make use of myelins biochemical characteristics (Kiernan, 2007). For example, LFB is likely attracted by the basic amino acids of myelins proteins and to a lesser extent also by phospholipids (Kiernan, 2007; Kluüver and Barrera, 1953; de Almeida and Pearse, 1958), whereas Gold atoms present in Gold stains are chemically reduced by myelin lipids in formaldehyde fixed tissue, leading to the black appearance of gold staining in myelin fibres (Schmued and Slikker, 1999; Schmued et al., 2008). While these stains are less specific to individual myelin proteins, their success is also sample-dependent: Vincze et al. (2008) found that in comparison to anti-MBP immunohistochemistry, LFB only revealed myelination in later developmental stages.

In summary, the variability in histological methods within our review highlights the need to use myelin histology specific for each experiment’s objective **(Recommendation 2**.**4)**. In contexts where myelin histology has never been used before, this might mean probing the specificity of different histological methods to the phenomenon of interest.

#### 4.4.4. Matching MRI and histology: ROI definition and image coregistration

Another factor that will affect MRI-histology correlations is the spatial coregistration between the two modalities. Even if both the microstructural MRI metric and the histology perfectly capture myelination, the correlation between the two will be low if the measures are taken from spatial locations that are not well matched between MR and histology. Here, the 2D nature of most histology techniques makes spatial coregistration challenging. Most of the assessed validation papers try to achieve spatial correspondence without coregistration (only 21 out of 70 studies use coregistration), but rather by manually outlining ROIs in the same anatomical location, identifying these ROIs based on landmarks.

The manual approach is commonplace, but has two pitfalls. First, placing ROIs may vary depending on the experimenter performing it, and automatic atlas-based ROI definitions are recommended practice in the neuroimaging field (Nichols et al., 2017). Second, coregistration may greatly improve the spatial correspondence between MR and histological images. While coregistration of MR images and histological images has unique challenges, such as the different spatial scales, changes in morphometry due to the tissue processing for histology, and potential cracks and folds in the tissue sections (Pichat et al., 2018), it has the advantage that ROI definition can be performed more reproducibly. Advances in automatic co-registration tools could aid a more widespread implementation in validation studies (Huszar et al., 2019) **(Recommendation 2**.**5)**.

### 4.5. Challenges in statistical comparisons of MRI and histology

Our meta-analysis focussed on studies using a correlative approach to measure the accuracy of MR metrics for validation. However, there are two key issues that are not addressed by correlative accuracy studies. First, validity depends on accuracy and accuracy depends on precision. Many of the studies in our review focus on accuracy, i.e. the measured parameter being in agreement with the true underlying biological parameter of interest. However, precision, i.e. low variability in repeated measurements, is also crucial to validity of a metric. In the case of MRI metrics, repetition across time (e.g. Arshad et al. (2017); Leévy et al. (2018)), across hardware (e.g. Bane et al. (2018); Leutritz et al. (2020)) and across sequences (e.g. Stikov et al. (2015a)) are all important, and, while outside the scope of this review, they are also a key prerequisite for accuracy.

Second, correlative approaches are key to assessing accuracy in histological validation studies, because for most MR metrics, there is no mathematical model capable of converting the units of the metric to readouts from myelin histology. However, the drawback of using shared variance as a measure for validity is that variance in the data is shaped by a variety of factors, including measurement noise and underlying variance in ‘ground-truth myelination’. Therefore, studies with high measurement noise or with low myelin variance (e.g. between subjects of the same group) may artificially deflate the true underlying correlation coefficient (Altman and Bland, 1983; Goodwin and Leech, 2006). Likewise, correlation does not equal causation and factors that affect both MRI and histology (such as post-mortem times) may artificially inflate or deflate correlation coefficients. A few studies have aimed to use non-correlative approaches for validation of MR metrics, for example by demonstrating effects from a given intervention on MR and histology (Lodygensky et al., 2012; Sampaio-Baptista et al., 2013), but fall outside the scope of this review.

#### 4.5.1. Achieving robust and meaningful correlations

As highlighted above, MR markers within our review have been validated with either within-subject or between-subject designs, and each design provides different information about the marker. We also found that a sizable subset of studies (n=19) pooled together multiple ROIs from multiple subjects, effectively measuring a mixture of both within and between-subject design, but without modelling each contribution independently. Additionally, in 8 studies, we could not deduce which variance was modeled. This limits the interpretability of the results, and fails to take the nested structure of the dataset into account. Therefore, future studies may want to perform analyses accounting for the subject structure in the data, thus maximising their interpretability **(Recommendation 3**.**1)**.

In studies measuring within-subject variance (i.e. spatial covariance of MR and histological metrics), we found two key drawbacks. First, correlation metrics are often reported for individual subjects, making it difficult to pool results across studies. Second, the extent of variability may not be comparable to between-subject variability, as contrast between GM and WM can be a strong driver of variance. This phenomenon is exemplified in Laule et al. (2006) and Peters et al. (2019), where correlation coefficients are lower in analyses including only ROIs from white matter, compared to analyses including a more heterogeneous set of ROIs. This is also reflected in our meta-analytical results for MTR (Figure 2, Supplementary 5), where the correlation coefficients were lower in between-subject studies (95% CI for *R*^2^: .15 and .51) compared to within-subject studies (95% CI for *R*^2^: .60 and .84).

In studies measuring between-subject variance a notable source of variability was that the data belonged to multiple groups (e.g. different ages, different pathology severity) (Thiessen et al., 2013). While this provides the correlation with more power, spurious correlations are a concern when dealing with multiple groups. Therefore, it is important for future validation studies to check test distributional assumptions, check the raw data for spurious correlations, and use correlation methods that are ro-bust to outliers (Wilcox, 2016; Salibian-Barrera and Zamar, 2002) **(Recommendation 3**.**2)**.

#### 4.5.2. Including potential biological confounds as co-variates

Myelin is not isolated from other microstructural features of brain tissue. In healthy tissue, myelin content is often related to iron and axon density. Iron occurs in high concentrations in oligodendrocytes and is important for the production and maintenance of myelin (Möller et al., 2019). It affects some MRI metrics similarly to myelin (Möller et al., 2019) and its distribution resembles myeloarchitecture (Fukunaga et al., 2010). Axonal density may also correlate with total amount of myelin, if axons are myelinated. This can be a problem for MR sequences that are sensitive to signals from axons, such as diffusion imaging, where axonal membranes affect diffusion signals even in the absence of myelin (Beaulieu, 2002). The concern about biological confounds is further corroborated by studies which perform MRI-histology correlations with various histological markers. For example, Jespersen et al. (2010) finds a correlation between FA and myelin stain of r =.78, and a correlation between FA and cell density of r = -.72, thus questioning the specificity of the FA-myelin correlation in the study.

For both iron and axons, colocalization can be even more pronounced in pathological samples. Conditions such as MS, or animal models such as Cuprizone-fed and Shiverer mice, are often used in validation studies, because of their known effect on myelin, but they also affect iron metabolism, due to iron’s role in myelin maintenance (e.g. Hametner et al. (2013); Pandur et al. (2019); Sergeant et al. (2005)) and axons, since myelin and axonal health are tightly coupled (Stassart et al., 2018).

Therefore, these biological confounds can often have an impact on the correlation coefficient between histology and MRI metrics. For example, an MRI marker sensitive to iron may yield a positive correlation with myelin histology, if myelin and iron colocalise in the tissue of interest. To test for myelin’s unique contribution to correlations, one needs to quantify and correct for these biological confounds **(Recommendation 3**.**3)**. Within the studies selected for our systematic review, only a few (n=7) performed iron histology, while half of the studies considered axons (n=35). Of these studies, only a small subset considered iron and axons as confounds in their analyses of myelin histology.

#### 4.5.3. Biases in the validation literature

Publication bias and questionable research practices have long been established as a key factor hindering pooling of evidence across studies in biomedical research (Ahmed et al., 2012). In our analyses, we find no evidence of abnormal *p*-value distributions in the validation literature, but we do find an asymmetry in the correlation between effect size and sample size of validation studies, which is often considered an indication of publication bias.

Asymmetric funnel plots can be driven by many factors. Selective outcome reporting and selective analysis reporting are the most common. However, not all asymmetric funnel plots are due to issues that impact meta-analytic inference (Sterne et al., 2011). For instance, in some circumstances sampling variation and chance can lead to an asymmetric funnel plot without real publication bias. Moreover, in the case of validation studies, it is expected that true effect sizes would be heterogeneous, especially when pooling together studies that examined how the same MR marker relates to different histological markers.

Our results show that correcting for publication bias would impact meta-analytic validation evidence for some of the MR metrics we analysed. These results need to be interpreted with caution, as our meta-analyses include relatively few studies, but are strengthened by the fact that 10 out of 70 studies in our review do not report results for all the metrics they collect. Taken together, these observations suggest that measures to prevent publication bias may be useful for future validation studies. Pre-registered protocols with pre-specified power analyses can strengthen the robustness of effect size estimates, and even provide more accurate estimates than meta-analytic analyses themselves (Kvarven et al., 2020). Therefore, using pre-registration may help establish more accurate effect sizes for the correlations between MR metrics and myelin histology **(Recommendation 3**.**4)**.

### 4.6. Conclusions and future directions

Our meta-analysis finds evidence for correlations between myelin histology and a range of MR markers: FA, RD, susceptibility, R1, MP and MTR, but not MD. These results verify that many MR markers are sensitive to myelin, but our analyses could not identify a single marker that is more sensitive to myelin than others. This suggests that for the time being, using multiple microstructural imaging markers in parallel may be the best way to test hypotheses related to myelin in humans in vivo.

We also find that the literature has a number of limitations. First, a wide variety of methodological approaches were used across the studies we assessed, making it challenging to estimate overall effect sizes and compare the effect sizes of different MR markers. Second, we find that heterogeneous and inconsistent reporting makes it difficult to assess the quality of the studies, and the factors driving differences in results. Third, we find some evidence of inflated effect sizes due to publication bias. Tackling these issues will be crucial to improving our strategies to measure myelin in humans.

## Acknowledgements

We thank Aman Badhwar for feedback on the meta-analysis. We thank Alex Bates, Michiel Cottar, Nicole Eichert, Heidi Johansen-Berg, Maria Morozova, Ruairi Roberts, Zeena Sanders and Nikolaus Weiskopf for feed-back on previous versions of the manuscript. We thank Edgar Liberis for his help with curating the online resource, Thomas Wassenaar for his advice on AMSTAR guidelines and Gunther Helms for his input on the macro-molecular pool measure. IL is funded by the Max-Planck-Society. AL is supported by a PhD Studentship from the Wellcome Trust (109062/Z/15/Z).

## Supplementary Figures and Tables

**Figure 1:**
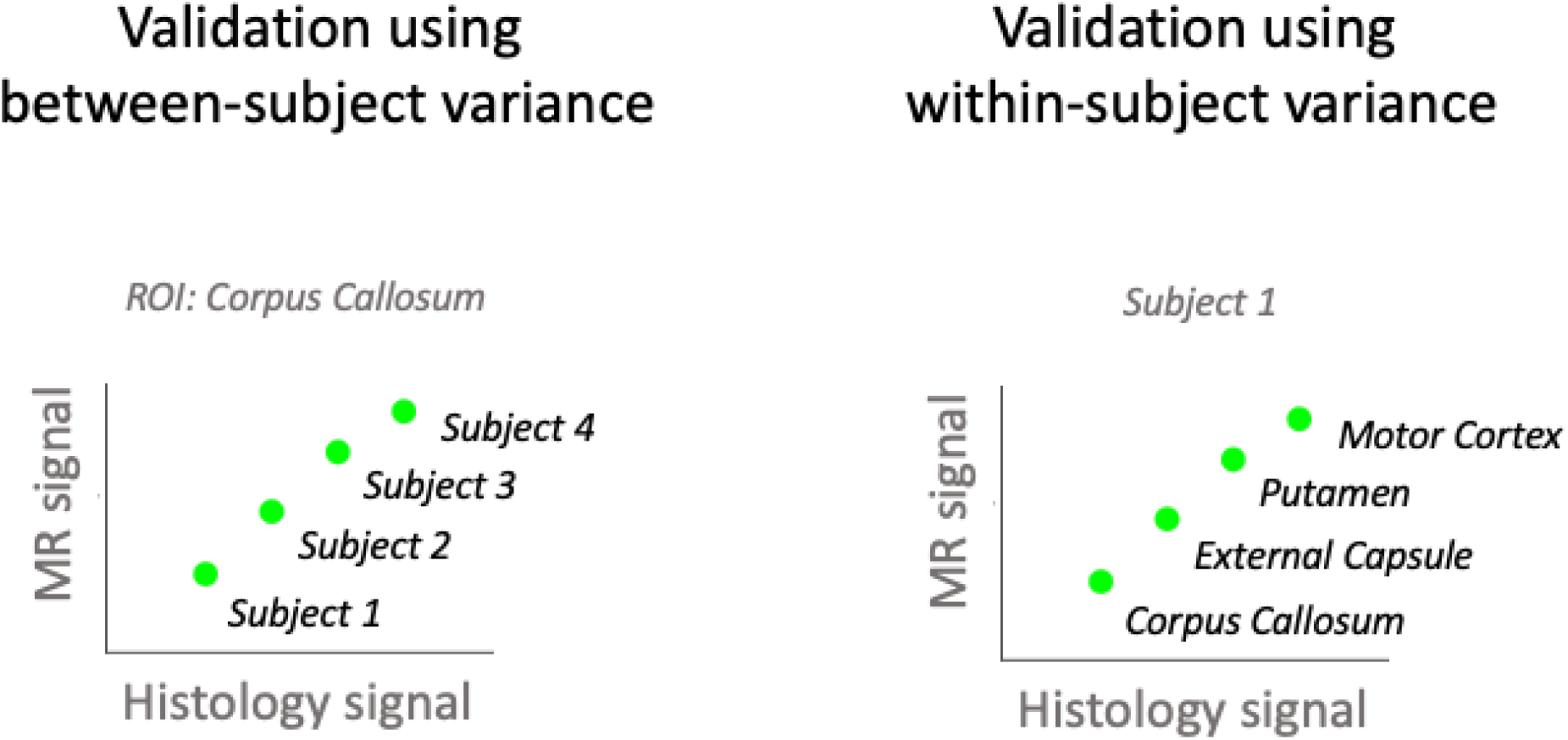
Mock examples of between-subject vs within-subject design in validation studies. In between-subject validation studies (left), each data point comes from a different subject and correlations are computed separately for each ROI. In within-subject validation studies (right), each data point comes from a different brain region and correlations are computed separately for each subject. In mixed designs, data points from different subjects and different regions are pooled into one data analysis.

**Figure 2:**
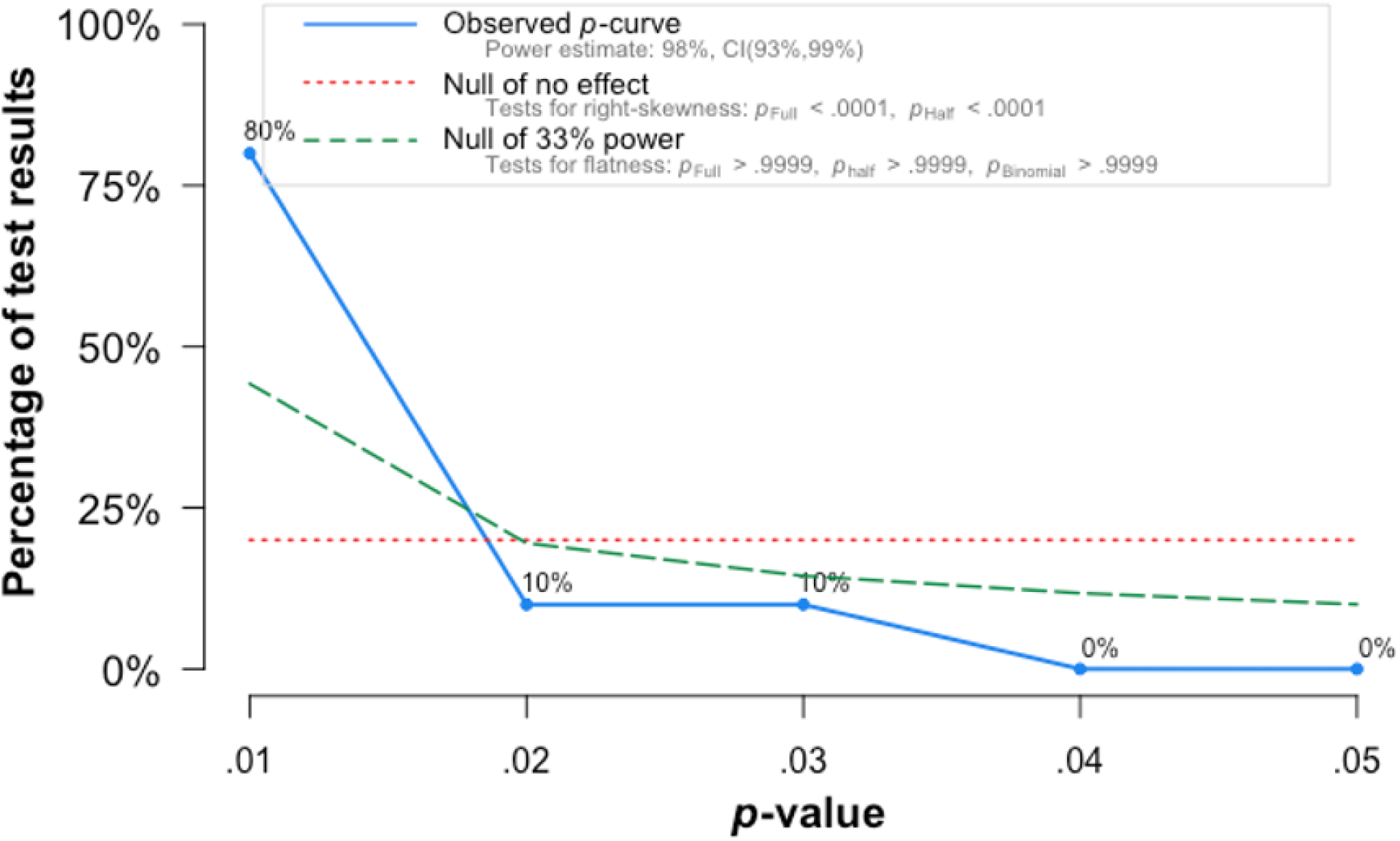
p-hacking in the validation literature. In fields where pressure to find significance biases published results p-value distributions tend to be right-skewed, i.e. to be characterized by p-values that are just below the 0.05 significance-threshold. We find evidence that the p-value distribution in the validation literature is, by contrast, left-skewed, with a majority of studies reporting p-values below 0.01.

**Figure 3:**
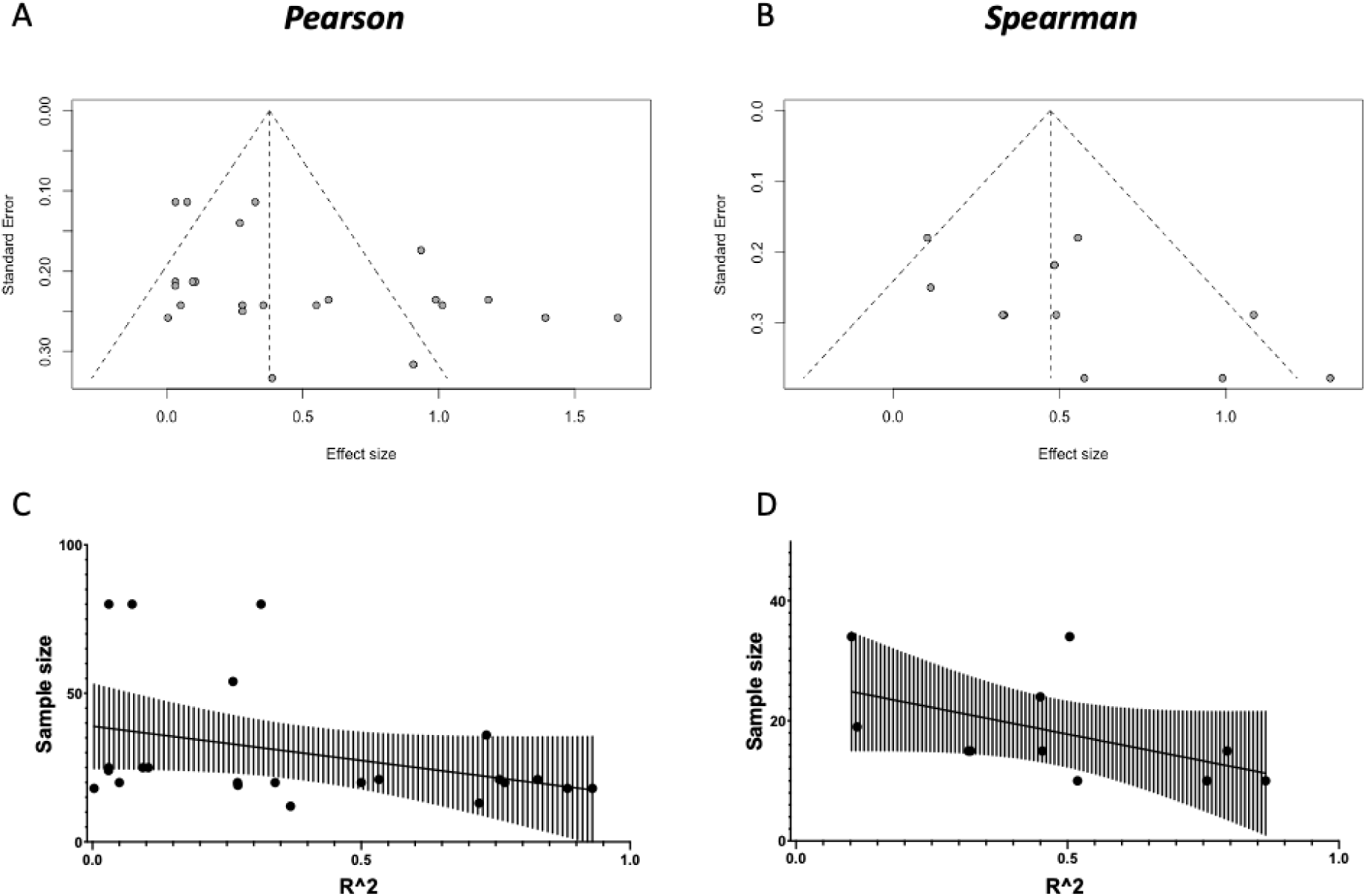
Publication bias. **[A and B]** Funnel plots of the studies in our quantitative meta-analyses (for Pearson and Spearman meta-analyses, respectively). We find some evidence for publication bias against low-sample size, low-effect articles (Eggers test: *t* = 2.419, *p* = 0.02471 for Pearson; *t* = 2.053, *p* = 0.07025 for Spearman). **[C and D]** To further characterize the relationship between Sample Size and Effect Size, we perform a correlation between the two, and find a relationship between higher sample sizes and lower effect sizes (Pearson: *r* = −0.3759, *p* = 0.0771; Spearman: *r* = −0.6451, *p* = 0.0365).

**Figure 4:**
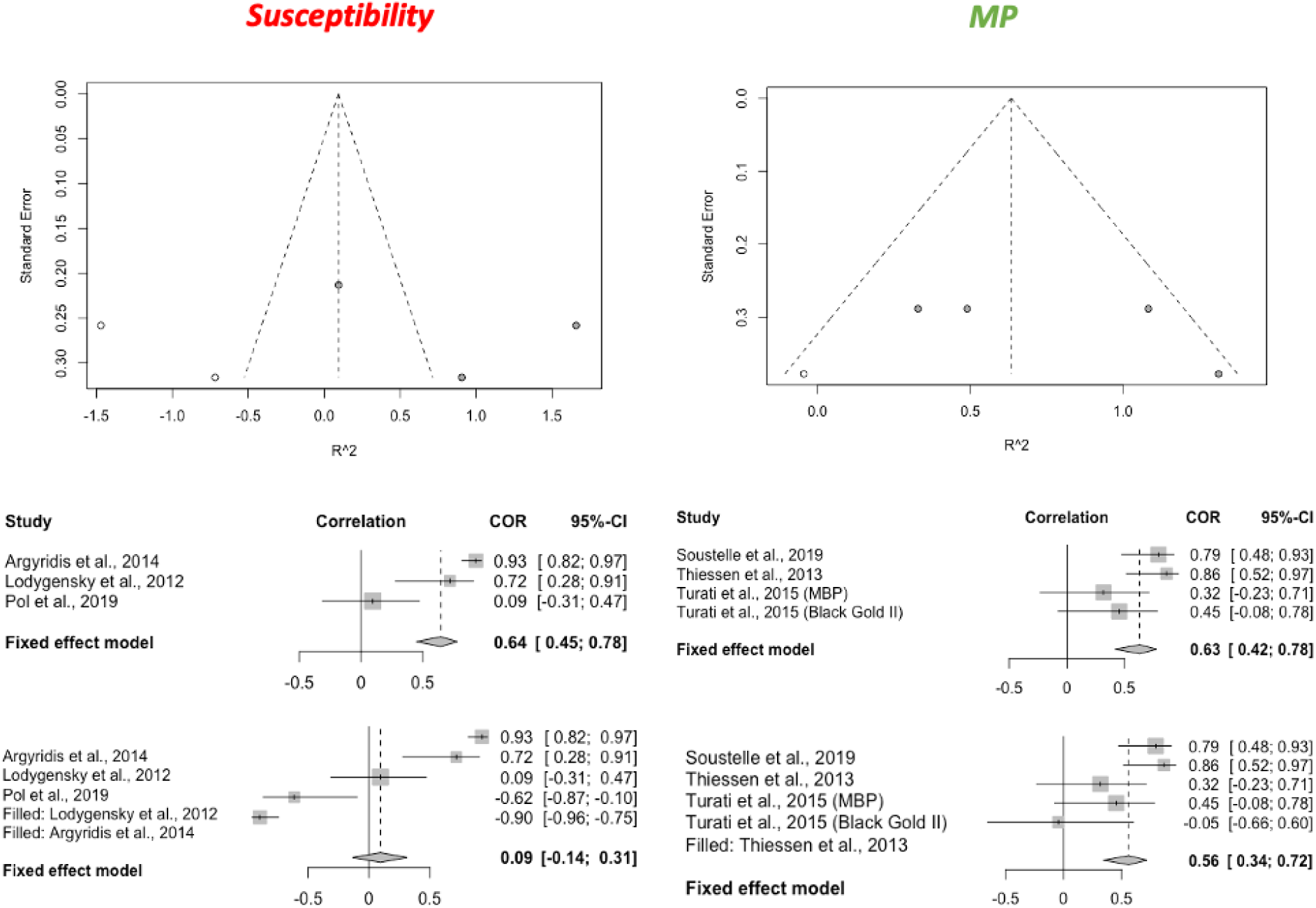
Trim-and-fill correction for publication bias. For both susceptibility and MP, we find evidence of publication bias (asymmetric funnel plots, top figures). We then use the trim-and-fill to correct for publication bias by imputing small-effect-small-sample studies that might have been performed, but not published. We then perform a meta-analysis on the imputed set of studies, and compare it to the complete case analysis in the main results.

**Figure 5:**
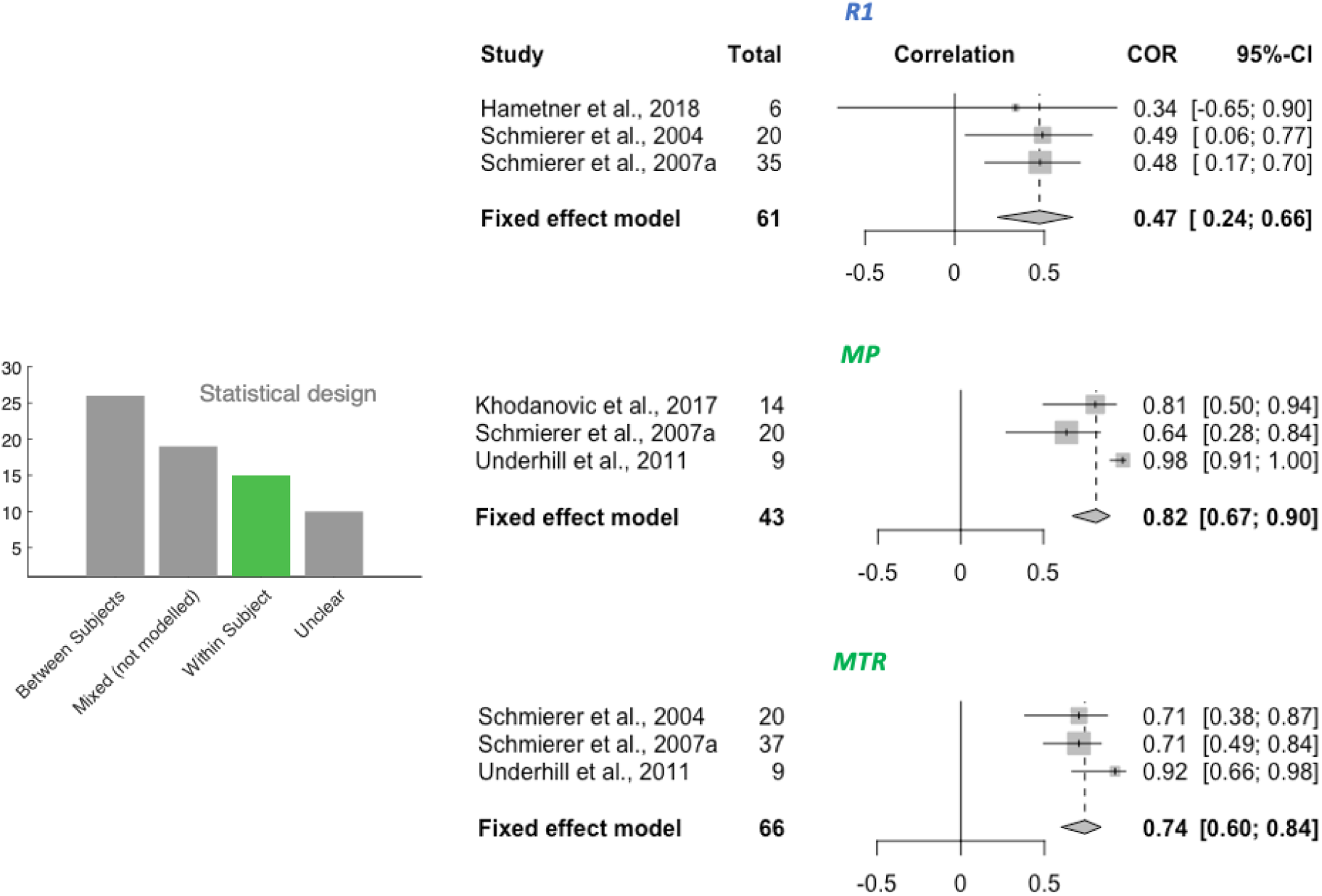
Within-subject meta-analyses. For the three metrics R1, MP and MTR, a meta-analysis on within-subject correlations could be performed. The resulting effect size estimate for MTR is higher than its between-subject effect size (see Figure 2).

**Table 1:**
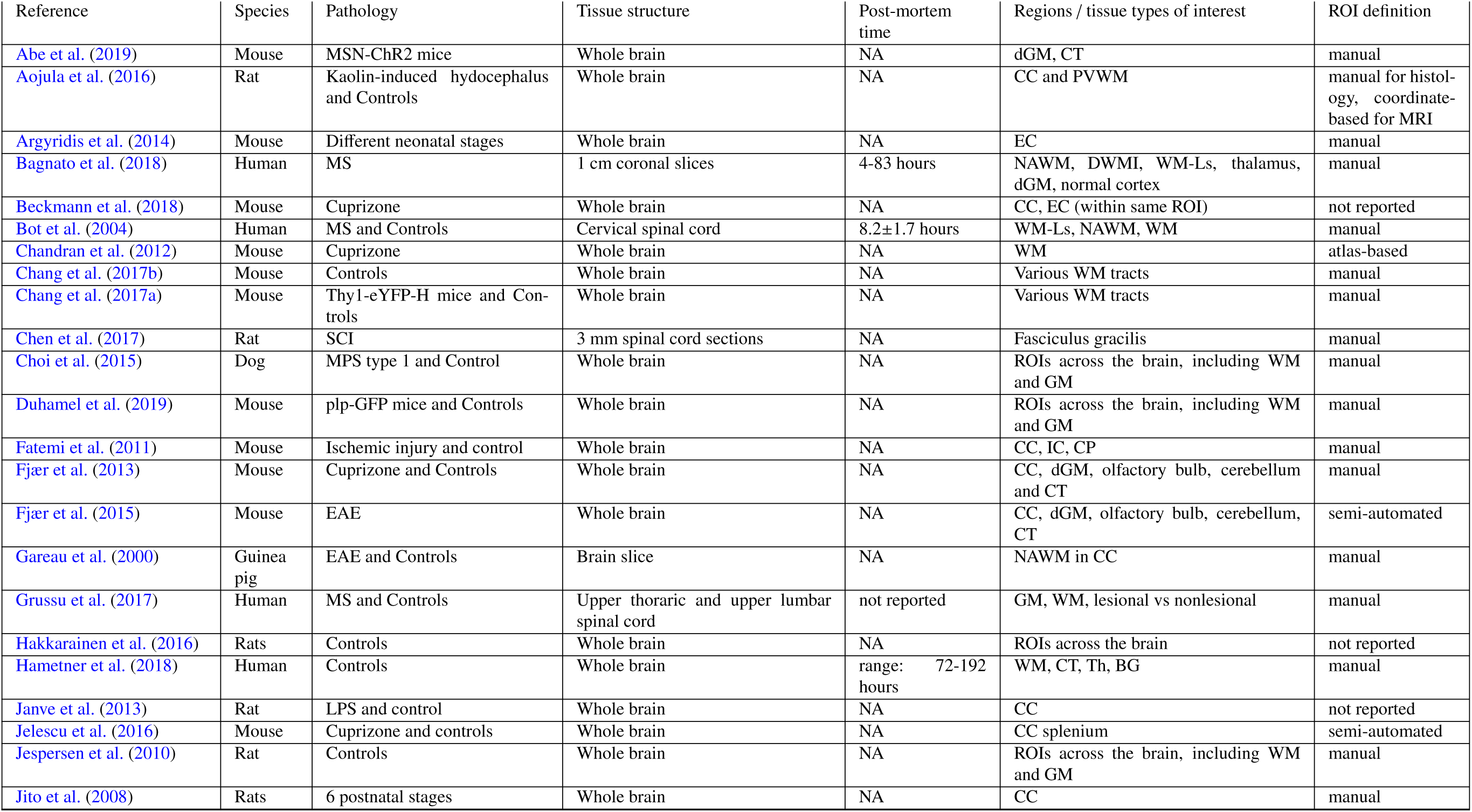
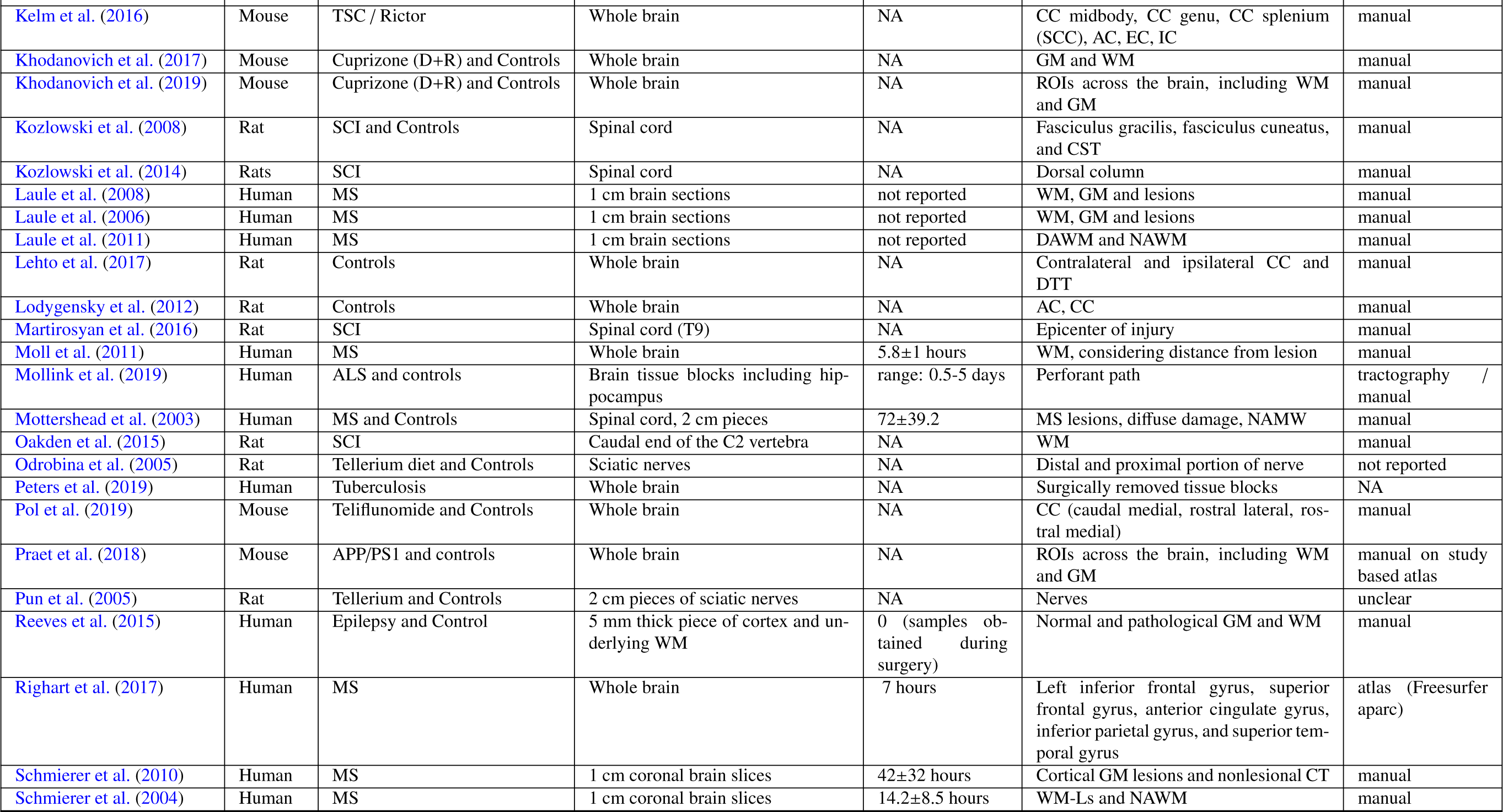
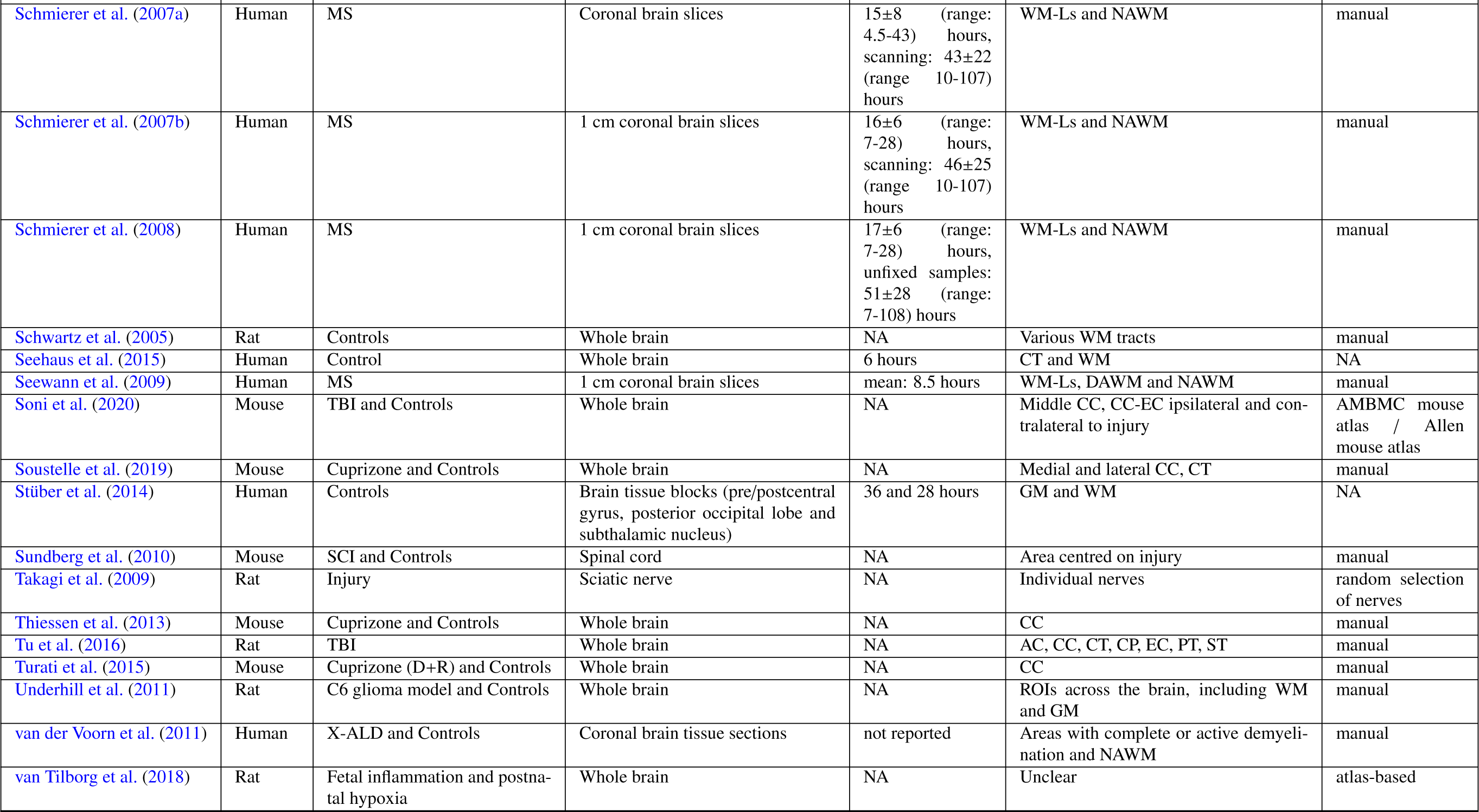
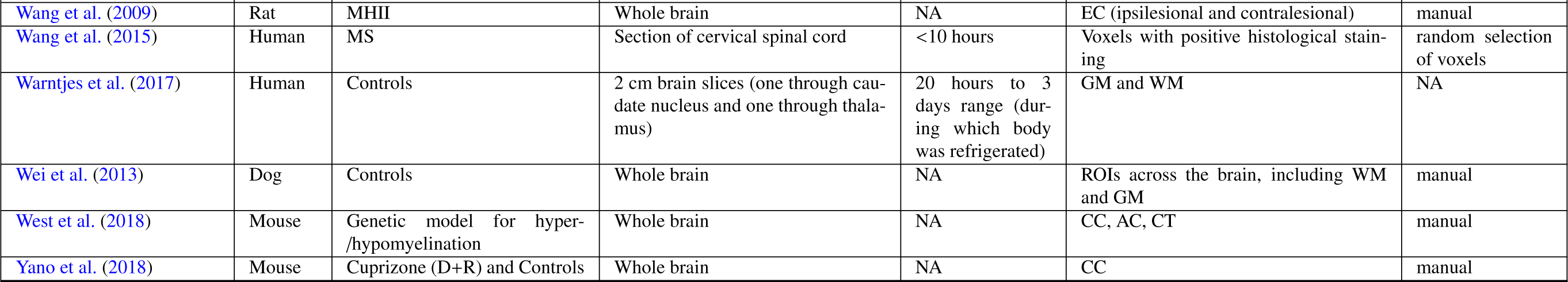
Basic information of the assessed validation studies. Provided are the species, condition/pathology studied, the tissue structure that underwent MR scanning, post-mortem time (if applicable), regions or tissue types of interest for the analysis and how they were defined. **Acronyms: ALS:** amytrophic lateral sclerosis; **APP/PS1:** Alzheimer mouse model; **Cuprizone:** Cuprizone-fed mice (D+R: demyelination and remyelination); **EAE:** allergic encephalomyelitis; **Kaolin:** Kaolin was used to induce communicating hydrocephalus; **LPS:** lipopolysaccharide-mediated animal model of MS, **MHII:** mild Hypoxic-Ischemic injury; **MPS:** mucopolysaccharidosis; **MS:** multiple sclerosis; **MSN-ChR2:** optical upregulation of Striatal medium spiny neurons **PLP-GFP:** Proteolipid protein - green fluorescent protein labelled mice; **TBI:** traumatic brain injury; **Thy1-eYFP-H:** mice that endogenously produce fluorescence signal; **TSC:** Tuberous sclerosis complex; **Shiverer:** Shiverer mice; **SCI:** spinal cord injury; **X-ALD:** X-Linked Adrenoleukodystrophy. **Anatomical structure: AC:** anterior commissure; **BG:** basal ganglia; **CC:** corpus callosum; **CP:** cerebellar peduncle; **CT:** cerebral cortex; **CST:** cortico-spinal tract; **DAWM:** diffusily abnormal white matter; **DTT:** dorsal tegmental tract; **DWMI:** diffuse white matter injury; **dGM:** deep gray matter; **GM:** gray matter; **NAWM:** normal appearing white matter; **OT:** optic tract; **PVWM:** Periventricular White Matter; **ST:** striatum; **Th:** Thalamus; **WM:** white matter; **WM-Ls:** white matter lesions.

**Table 2:**
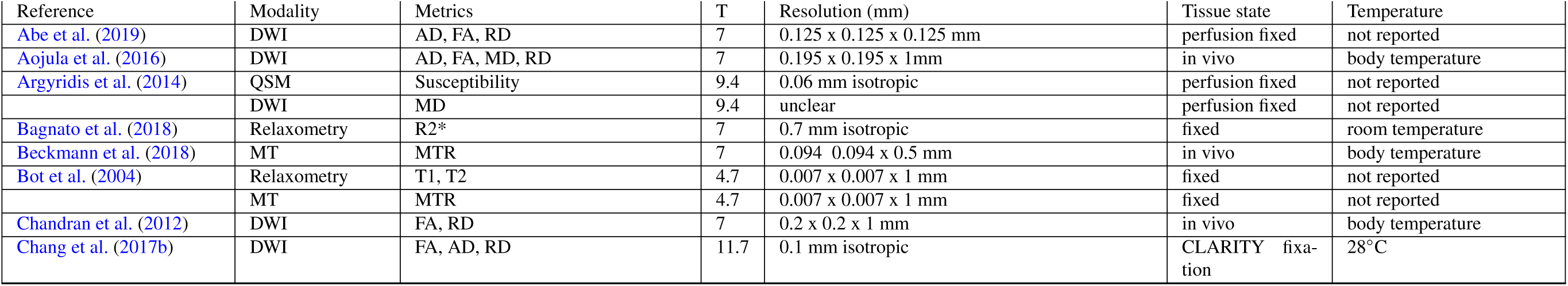
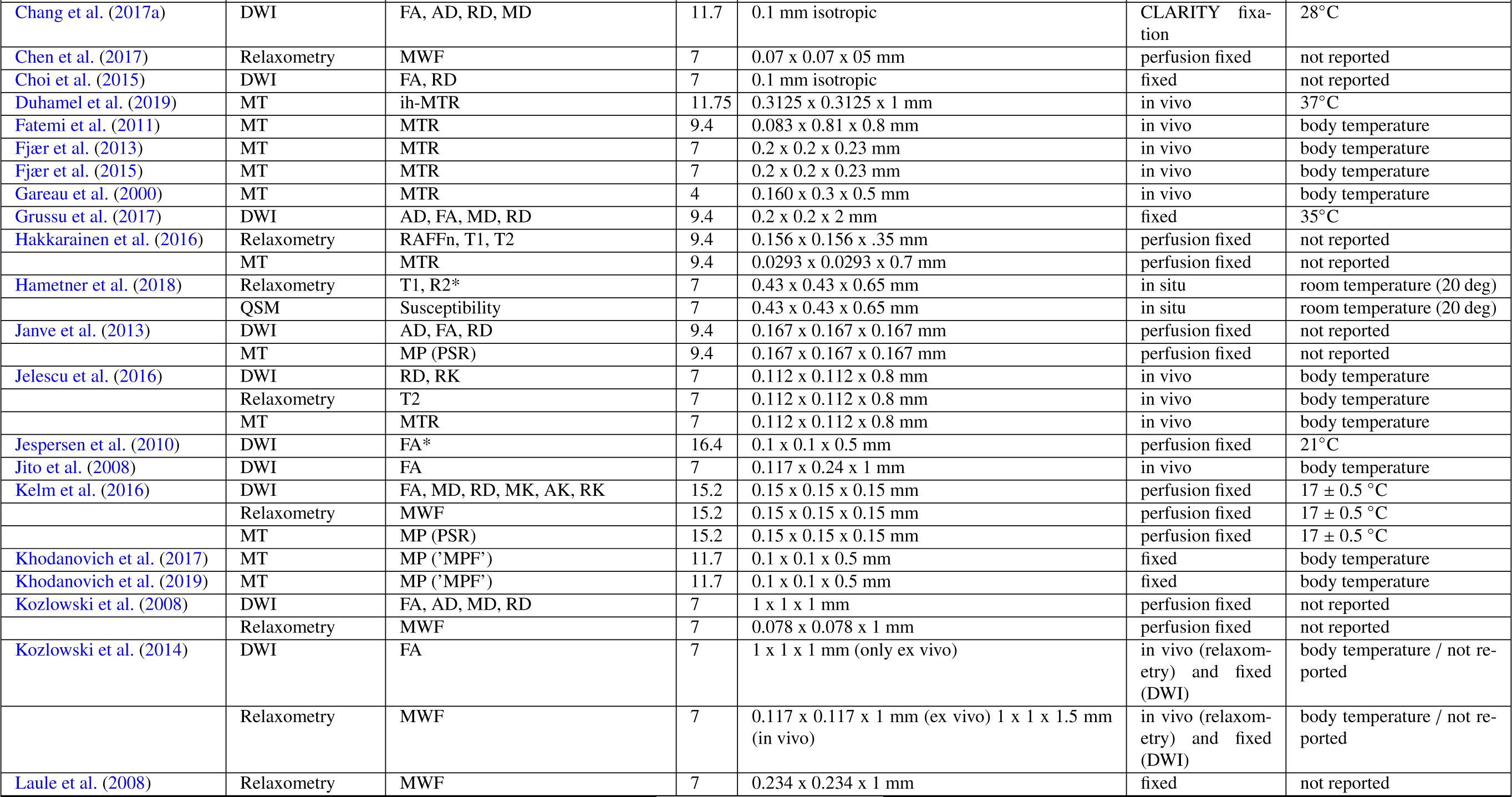
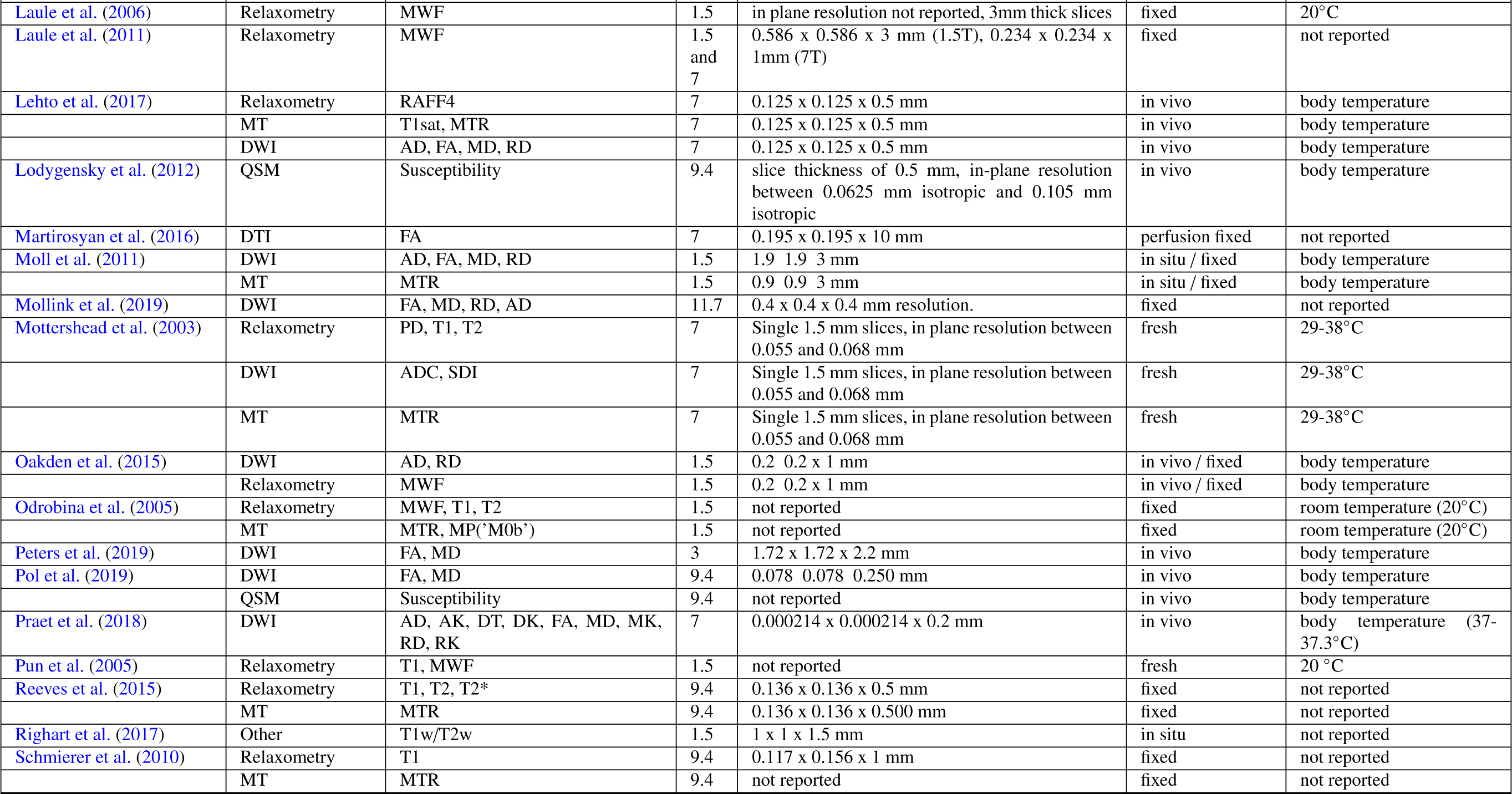
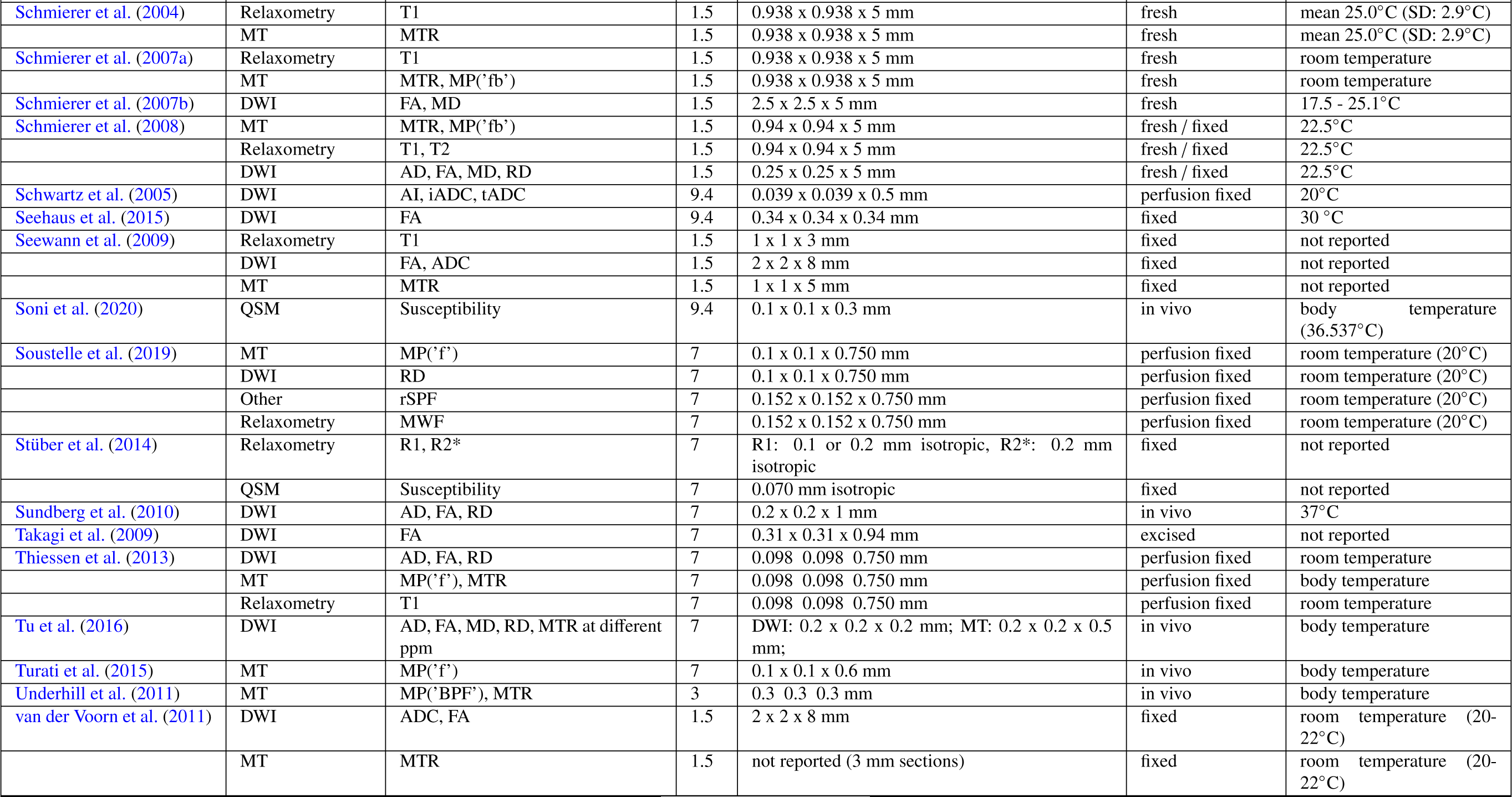
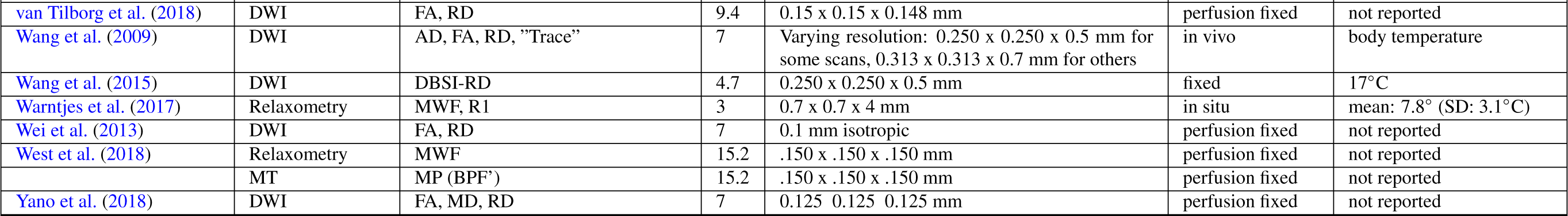
Information on MRI imaging of the assessed validation studies. Provided are imaging modality, field strength in Tesla (T), imaging resolution in terms of voxel size, and the tissue state and temperature during scanning. **Acronyms: Imaging: DWI: Di***ff***usion-weighted imaging**: **AK**: axial kurtosis; **AI**: Anisotropy Index (tADC/lADC); **DBSI-RD**: diffusion basis spectrum imaging - based radial diffusivity; **DK**: diffusion kurtosis metrics; **DT**: diffusion tensor metrics; **FA**: fractional anisotropy (from diffusion tensor model); **lADC**: longitudinal apparent diffusion coefficient (not modelled with tensor); **MD**: mean diffusivity (from diffusion tensor model); **MK**: mean kurtosis; **RD**: radial / transverse diffusivity (from diffusion tensor model); **RK**: radial kurtosis; **SDI**: diffusion standard deviation index; **tADC**: transverse apparent diffusion coefficient (not modelled with tensor); **Relaxometry**: **MWF**: myelin water fraction; **R1**: longitudinal relaxation rate; **R2\***: effective transverse relaxation rate; **RAFF4**: Relaxation Along a Fictitious Field in the rotating frame of rank 4; **T1**: longitudinal relaxation time; **T2**: transverse relaxation time; **T2\***: effective transverse relaxation time; **MT: magnetisation transfer**: **BPF**: bound pool fraction; **F**: pool size ratio; **Fb**: macromolecular proton fraction; **ih-MTR**: MTR from inhomogeneous MT; **M0b**: fraction of magnetization that resides in the semi-solid pool and undergoes MT exchange; **MP**: macromolecular pool; **MPF**: macromolecular proton fraction; **MTR**: magnetisation transfer ratio; **PSR**: Macromolecular-to-free-water pool-size-ratio; **STE-MT**: MTR based on short echo time imaging; T1sat: T1 of saturated pool; **UTE-MTR**: MTR based on ultrashort echo time imaging; **QSM**: quantitative susceptibility mapping; **Others: rSPF**: relative semi-solid proton fraction from an 3D ultrashort echo time (UTE) sequence within an appropriate water suppression condition; **T1w**/**T2w**: ratio of image intensity in a T1-weighted vs T2-weighted acquisition.

**Table 3:**
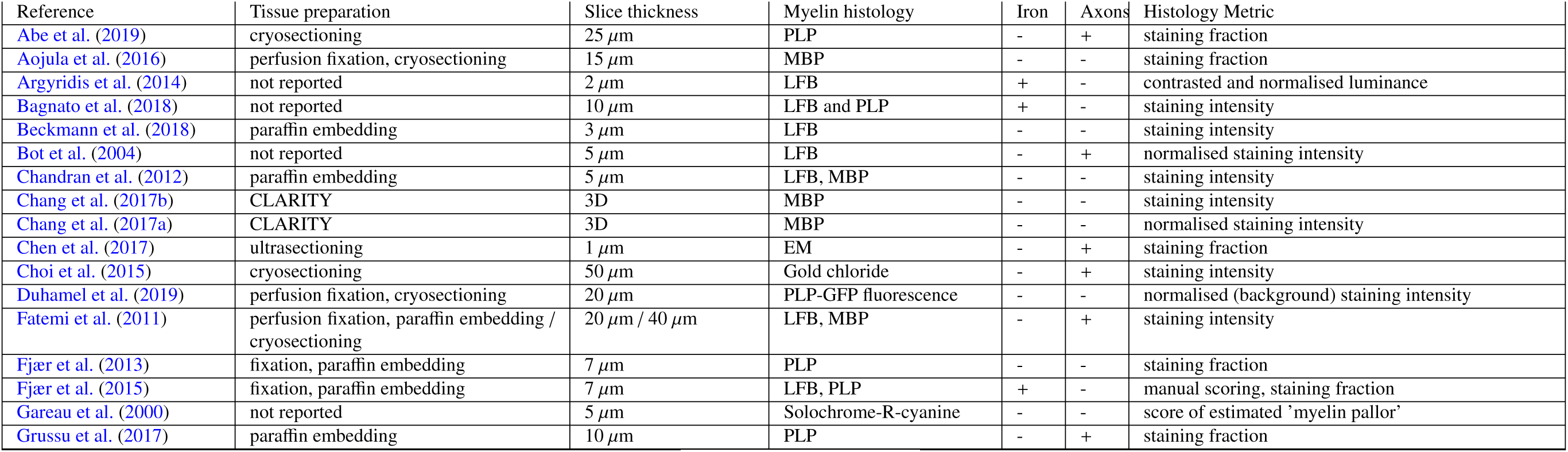
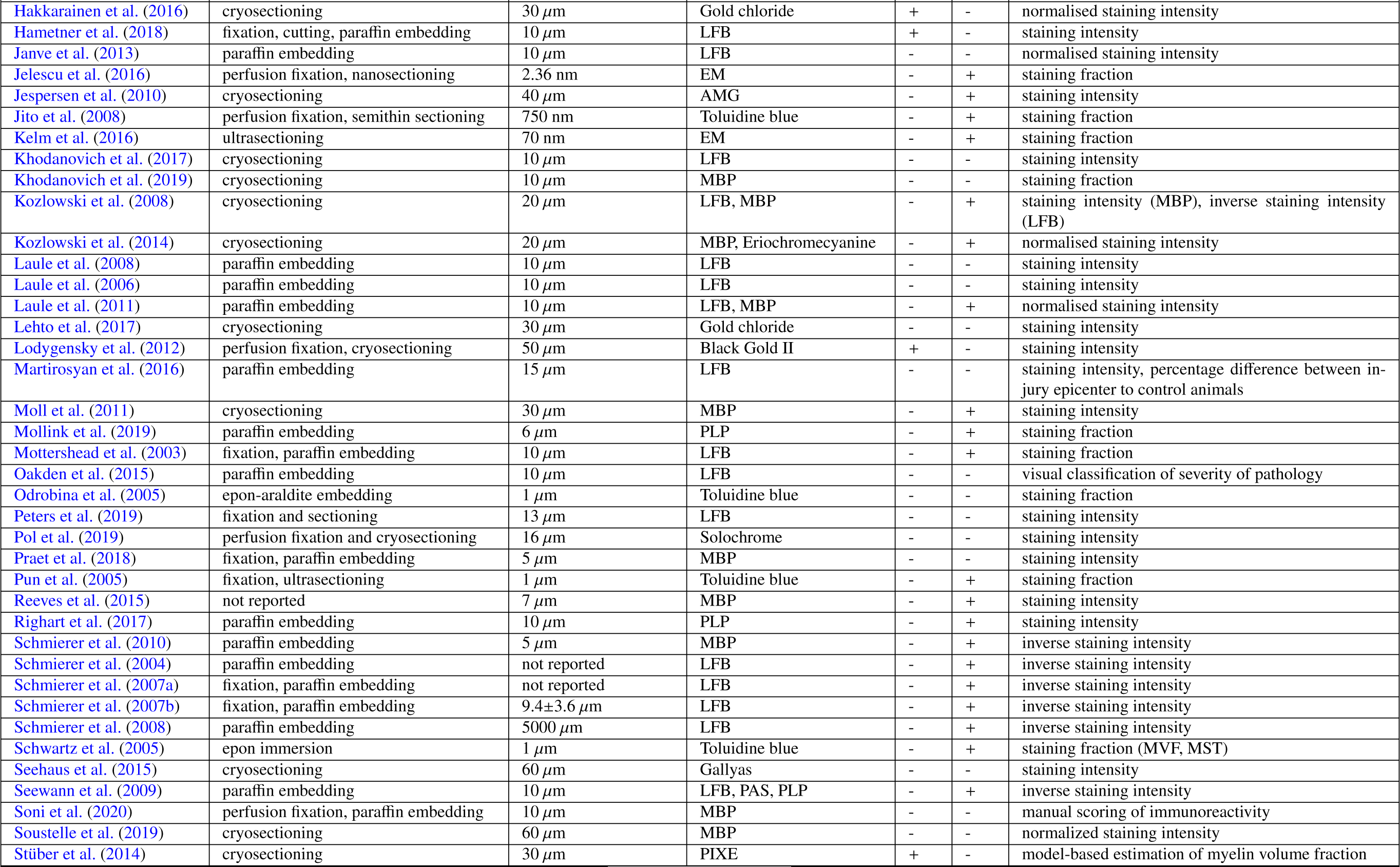
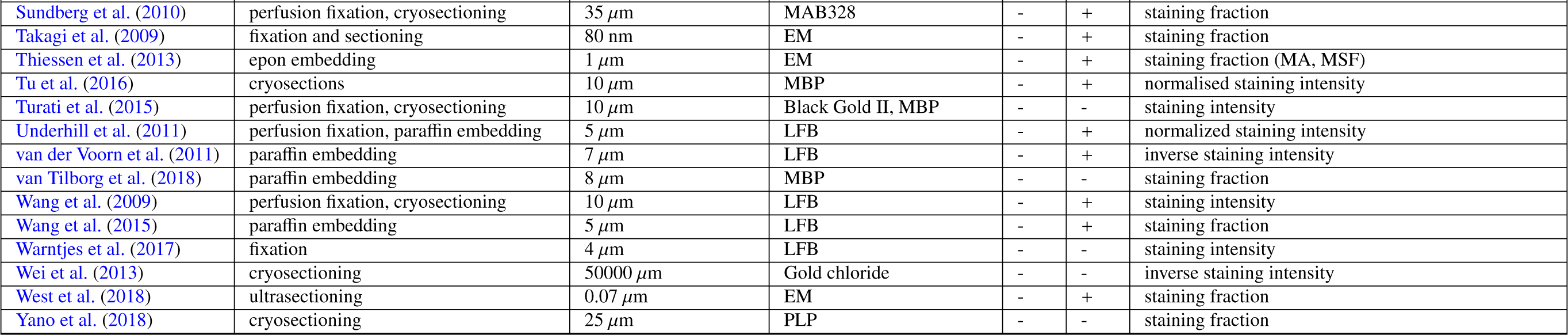
Information on histology of the assessed validation studies. Provided are information in tissue preparation, histological slice thickness, the histological method for myelin (and whether iron and axons were also considered) and the obtained histological metric. **Acronyms: AMG**: Autometallographic myelin stain; **LFB**: Luxol fast blue stain; **MA**: fraction of myelinated axons; **MAB238**: Anti-oligodendrocyte immunohistochemistry; **MBP**: Anti-myelin-basic-protein immunohistochemistry; **MSF**: myelin sheath fraction; **MST**: myelin sheath thickness; **MVF**: myelin volume fraction; **PAS**: periodic acid-Schiff; **PIXE**: proton-induced X-ray emission; **PLP**: Anti-proteolipid-protein immunohistochemistry.

**Table 4:**
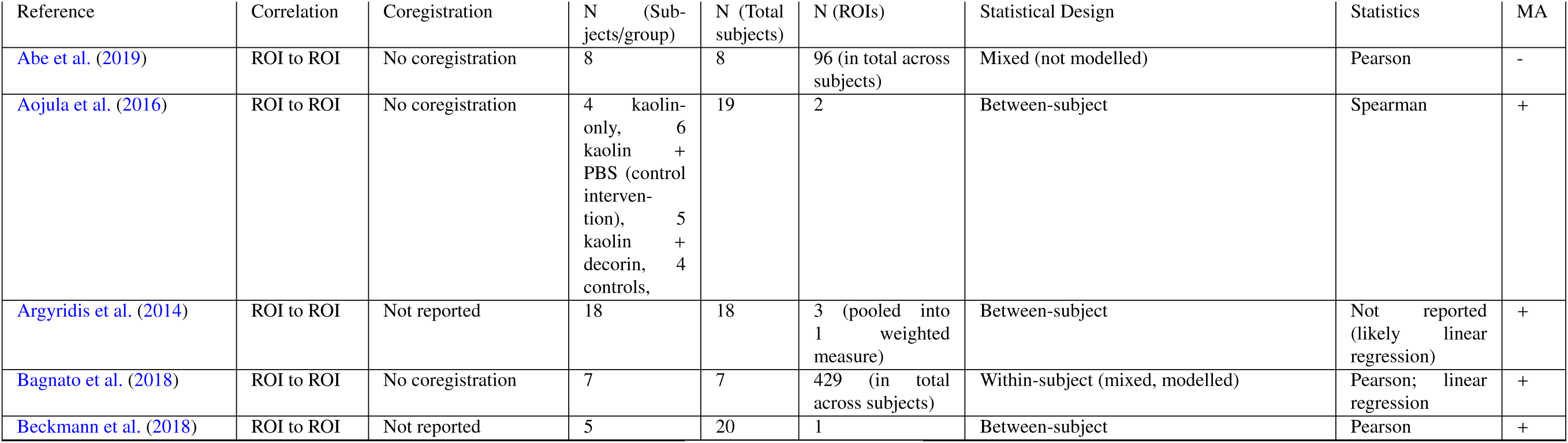
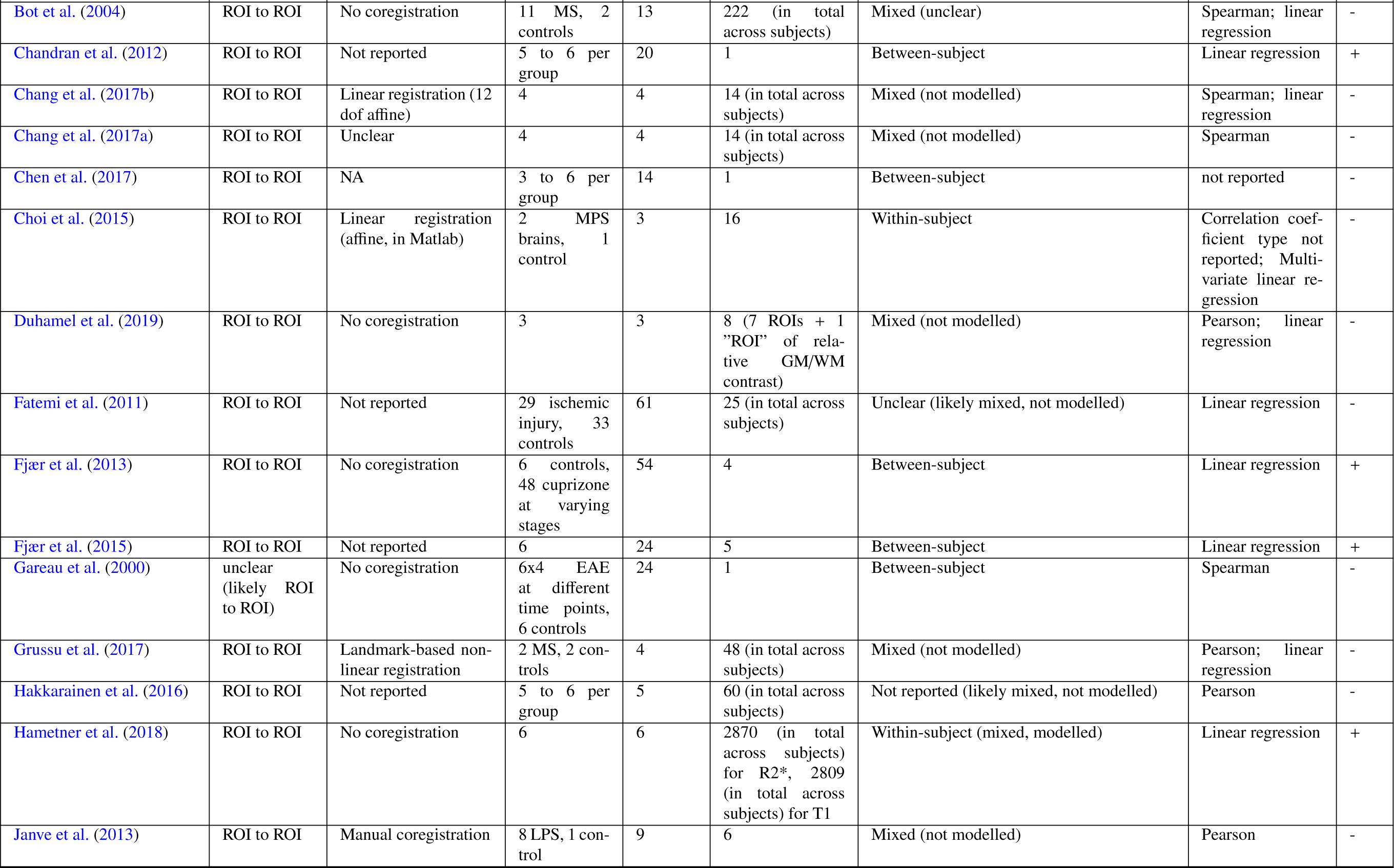
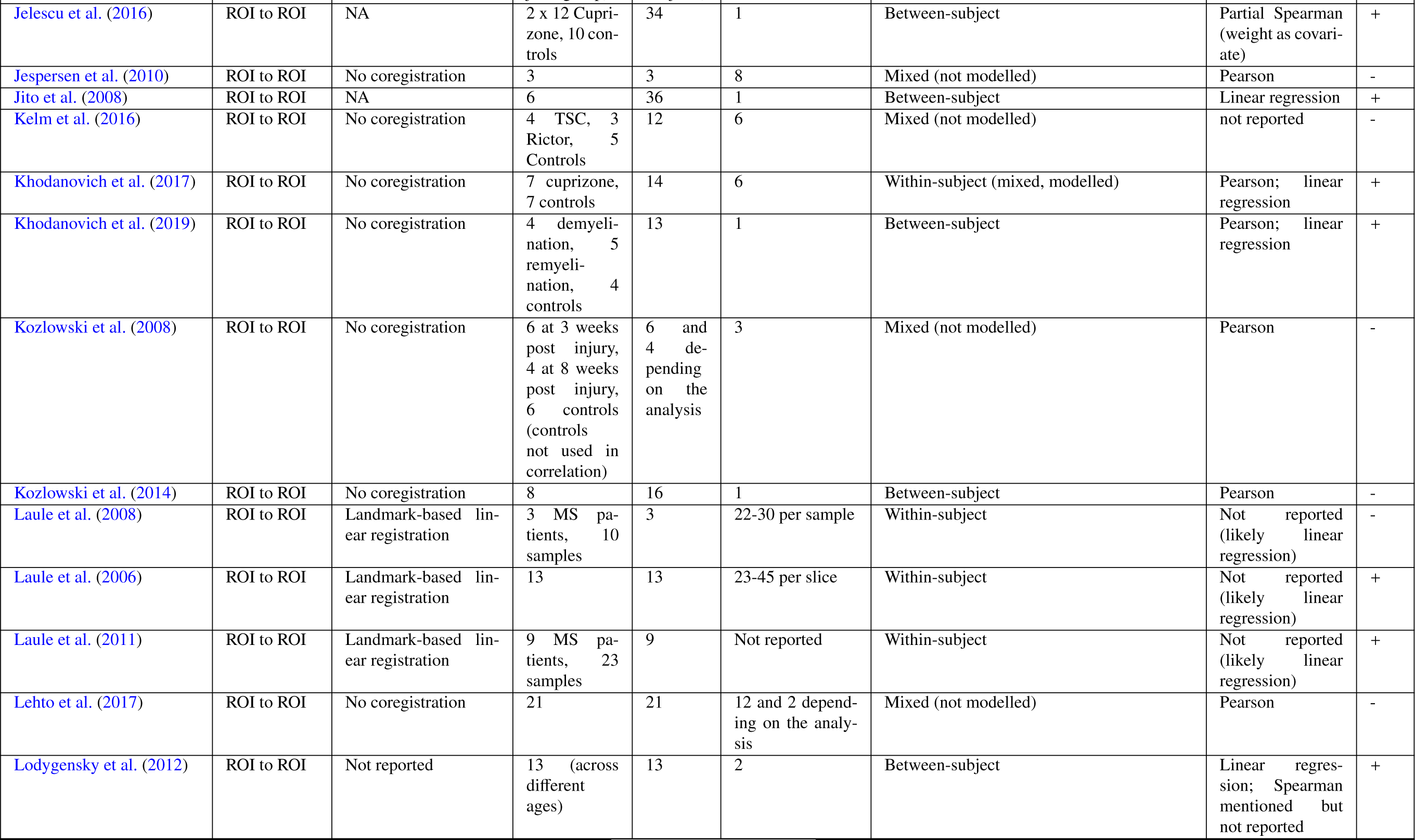
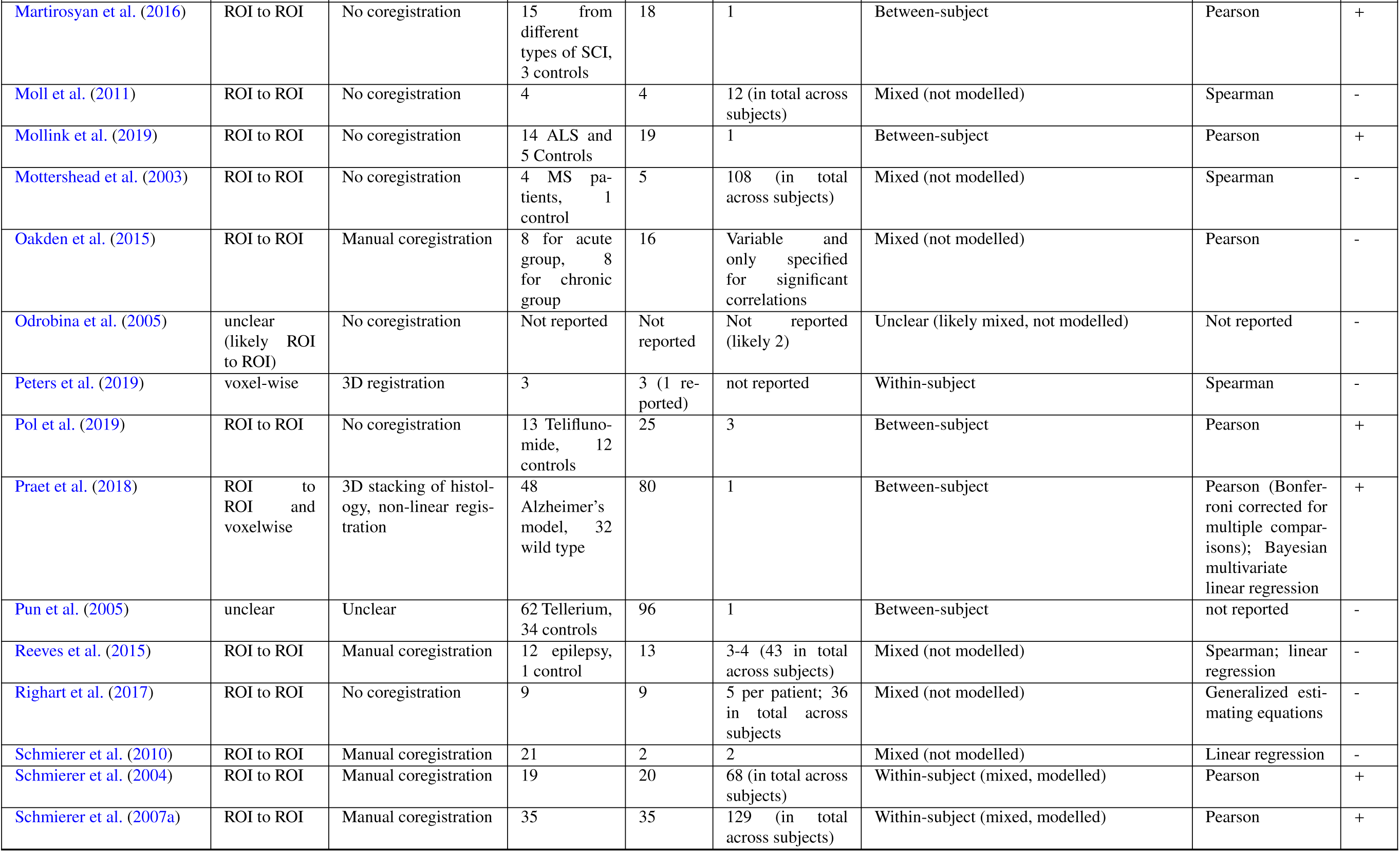
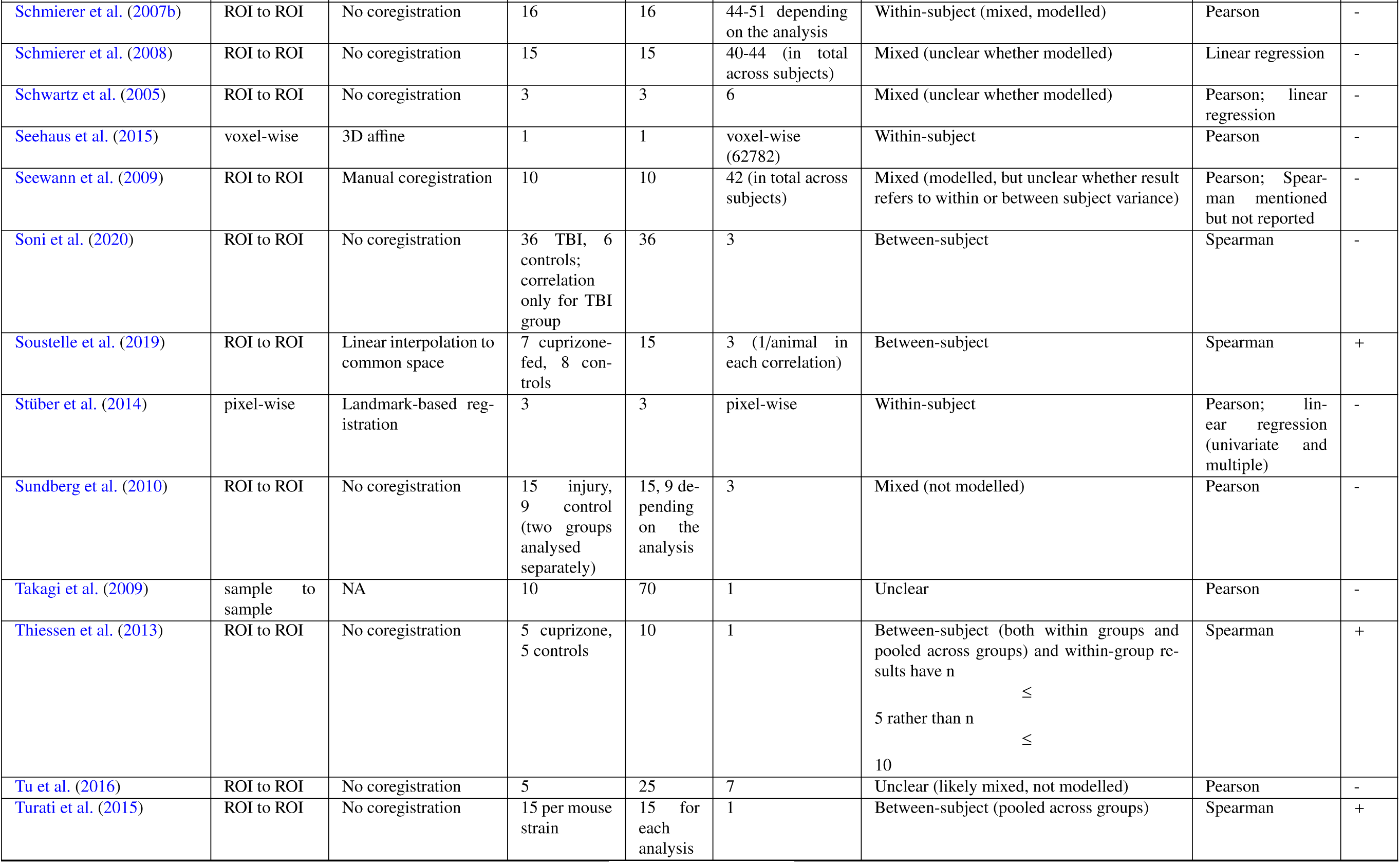
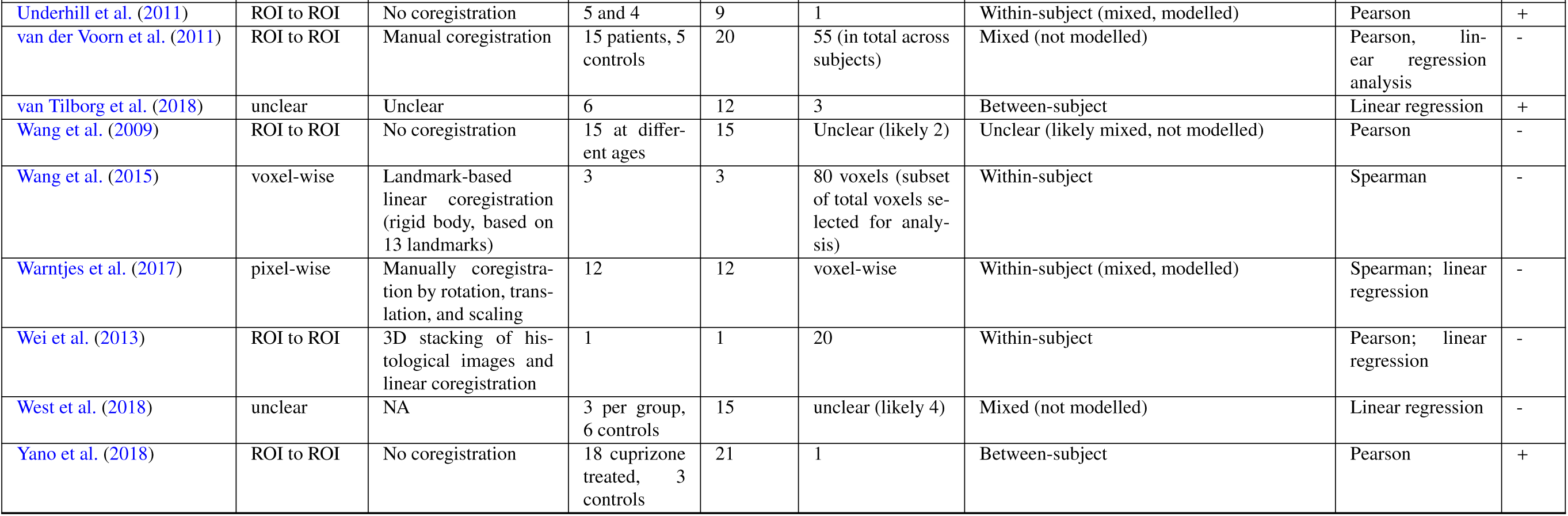
Information on the employed statistical approach of the assessed validation studies. Provided are information on the nature of data points that entered the correlation analysis, the method for coregistration between MRI and histology, sample size (number of subjects per group, total number of subjects, and number of data points for the analysis), the type of variance that was modelled in the statistical analysis, the reported statistic, and whether the study was selected for our meta-analysis (MA).

**Table 5:**
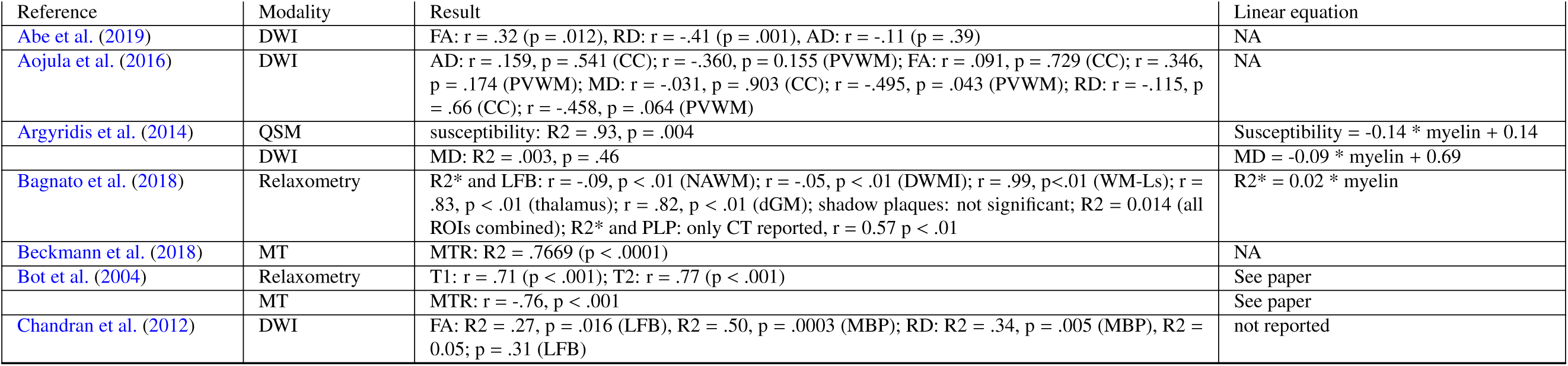
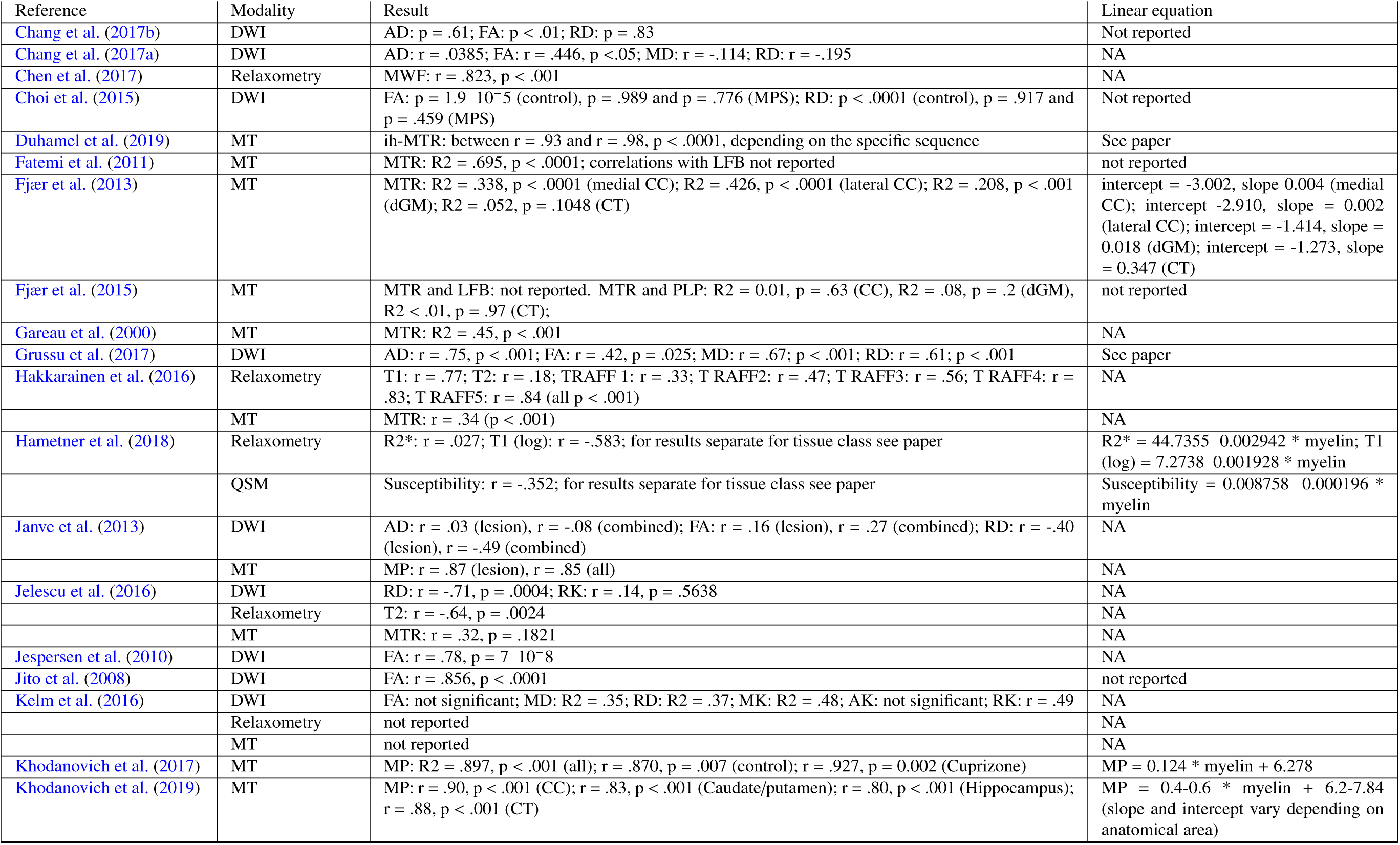
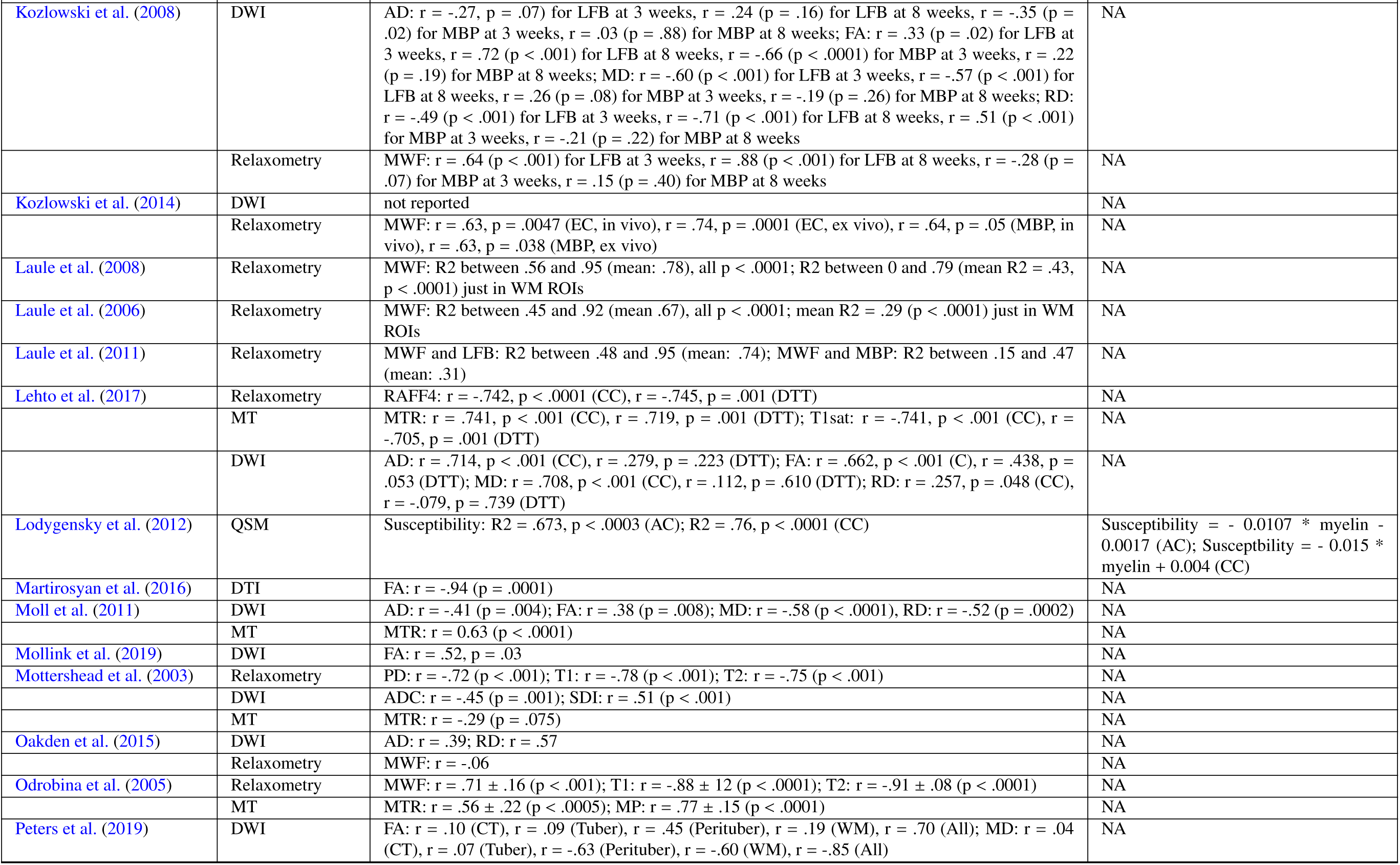
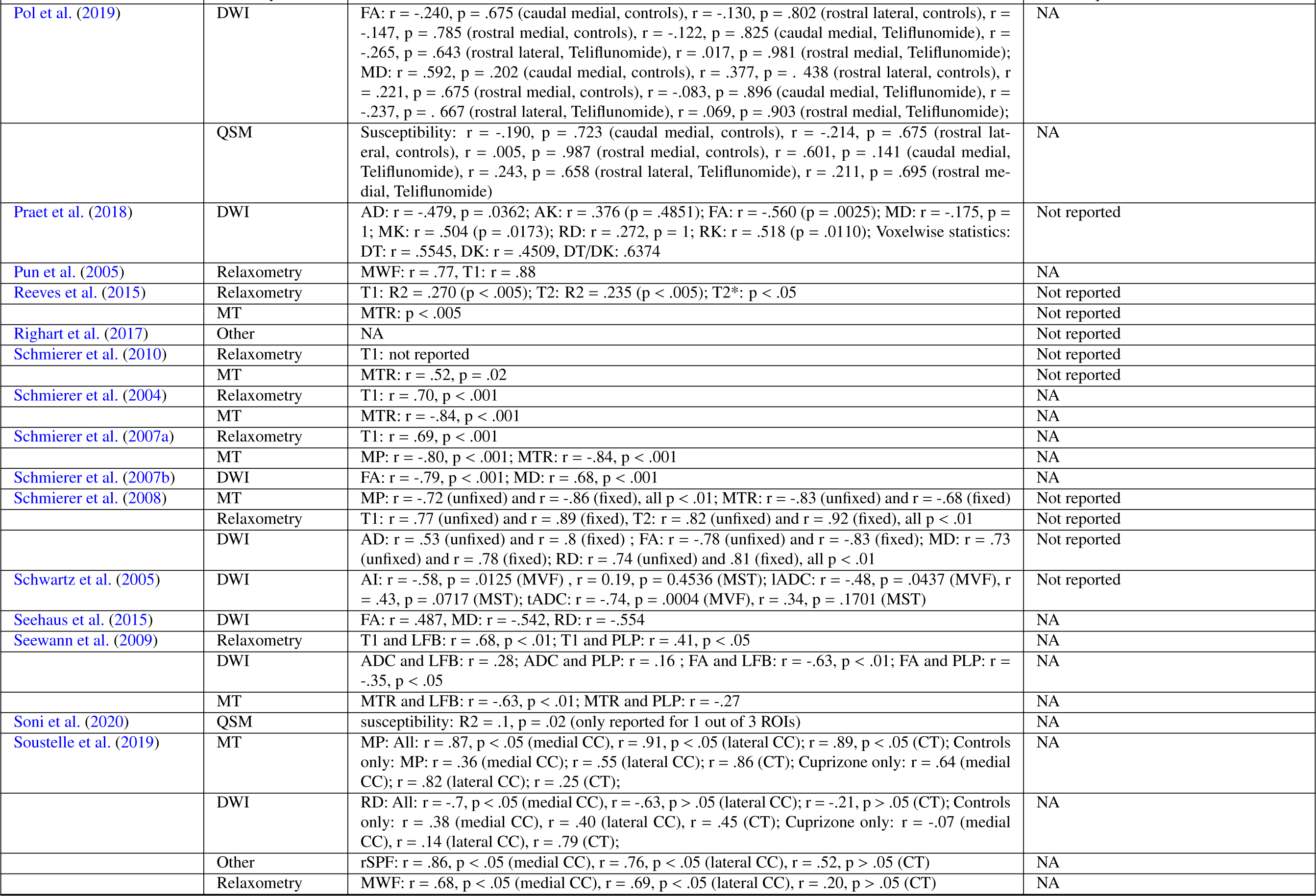
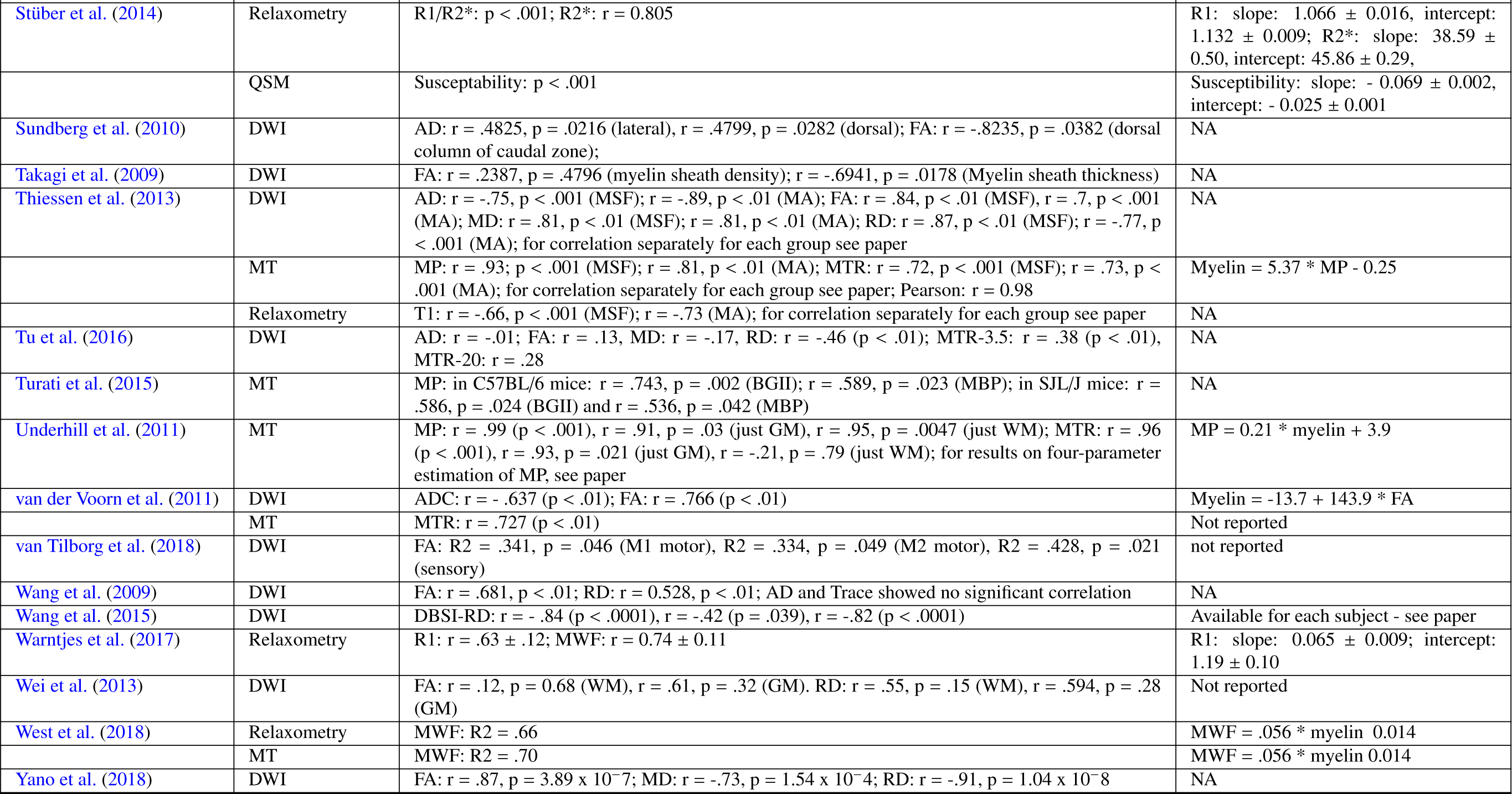
Information on the statistical results of the assessed validation studies.

